# Unify: Learning Cellular Evolution with Universal Multimodal Embeddings

**DOI:** 10.1101/2025.09.07.674681

**Authors:** Huawen Zhong, Wenkai Han, Guoxin Cui, David Gomez Cabrero, Jesper Tegner, Xin Gao, Manuel Aranda

**Affiliations:** BioEngineering Program, Biological and Environmental Science and Engineering Division, King Abdullah University of Science and Technology (KAUST), Thuwal, 23955-6900, Kingdom of Saudi Arabia; Klarman Cell Observatory, Broad Institute of MIT and Harvard, Cambridge, MA 02142, United States; Marine Science Program, Biological and Environmental Science and Engineering Division, King Abdullah University of Science and Technology (KAUST), Thuwal, 23955-6900, Kingdom of Saudi Arabia; Computer Science Program, Computer, Electrical and Mathematical Sciences and Engineering Division, King Abdullah University of Science and Technology (KAUST), Thuwal 23955-6900, Kingdom of Saudi Arabia; Unit of Computational Medicine, Department of Medicine, Center for Molecular Medicine, Karolinska Institute, Karolinska University Hospital, Stockholm, Sweden; Science for Life Laboratory, Solna, Sweden; Center of Excellence on Smart Health, King Abdullah University of Science and Technology (KAUST), Thuwal 23955-6900, Kingdom of Saudi Arabia; Center of Excellence for Generative AI, King Abdullah University of Science and Technology (KAUST), Thuwal 23955-6900, Kingdom of Saudi Arabia

**Author notes:** Corresponding author: Xin Gao, Manuel Aranda. The first two authors should be regarded as joint First Authors.

**Keywords:** Cross-species integration, scRNA-seq, cell type evolution, perturbation

## Abstract

Integrating single-cell RNA-sequencing (scRNA-seq) data across species is hindered by evolutionary divergence, technical batch effects, and the reliance on one-to-one orthologs. We present Unify, a transfer learning methodology that learns universal cell embeddings by defining functionally coherent, multi-modal macrogenes. This is achieved by combining RNA expression with embeddings from protein language models and general-purpose language models. Unify transcends species boundaries, enabling cross-species comparisons beyond strict gene-level homology. Unify corrects batch effects while preserving conserved biological signals across vast evolutionary distances and enables more accurate prediction of perturbation responses across species, such as from mouse to human. Applied to species separated by over 700 million years, Unify reconstructs more accurate multi-species cell-type evolutionary trees and uncovers convergent gene programs. Together, these results establish Unify as a powerful method for comparative single-cell genomics and evolutionary biology.

## Introduction

The advent of single-cell RNA-sequencing (scRNA-seq) has revolutionized our ability to map the landscape of cellular identity and function within individual organisms^1-3^. An exciting and critical new frontier is comparative single-cell genomics, which seeks to extend these insights across the tree of life. By comparing cell atlases from different species, we can uncover the fundamental principles of gene regulation that are conserved through evolution, trace the origins and diversification of cell types, and critically, enhance the translation of biological findings from model organisms to human health^3-6^. However, to achieve meaningful cross-species integration of single-cell data is a significant challenge. The task is profoundly challenging due to the need to distinguish conserved biological signals from the confounding effects of technical variability between experiments and, more significantly, the deep evolutionary divergence that has reshaped genes and genomes over millions of years^7-9^.

There is a growing interest in cross-species data integration, and several methods have been developed to address this challenge^6, 10-13^. Early computational efforts relied heavily on one-to-one orthologs as anchors to map transcriptomes across species^14-20^. More recently, methods like SAMap^6^ and CAME^13^ have advanced this by utilizing BLAST-based homology to construct cross-species graphs. Furthermore, a newer generation of approaches, including SATURN^10^, UCE^11^, and TranscriptFormer^12^, leverage embeddings from protein language models like ESM-2^21^ to capture functional relationships between both orthologous and non-orthologous genes. However, their reliance on a single modality, i.e. protein sequence, provides an incomplete picture of gene function, neglecting the rich, contextual information captured in curated, expert-annotated biological knowledge bases. While text-based foundation models, such as GPT-4^22^ and Gemini-2.5^23^, have shown promise in encoding this knowledge for gene annotation, they have not yet been integrated into cross-species cell embedding frameworks. Furthermore, a difficult trade-off exists for current methods. Aggressive batch correction often comes at the cost of losing subtle but critical biological information across species^7-9^. Additionally, existing integration approaches typically produce either corrected expression profiles restricted to one-to-one orthologous genes^14-17^ or integrated embeddings^6, 10, 18-20^ that lack direct interpretability. Currently, no method excels at both robust batch correction and faithful preservation of conserved biology, thereby limiting sensitive downstream applications, such as cross-species perturbation prediction or the reconstruction of cell-type phylogenies^8, 9^.

Here, we introduce Unify (<u>U</u>niversal <u>N</u>on-orthologous Integration via <u>F</u>unctional <u>Y</u>ield), a transfer learning framework designed to resolve the core challenges of cross-species single-cell analysis. Unify builds on the concept of functionally coherent macrogenes that transcend species boundaries, extending it with a multimodal representation. While conceptually related to approaches using protein embeddings (e.g., SATURN), Unify uniquely integrates structural information from protein language models with functional context derived from general-purpose large language models interpreting biological knowledge. To achieve this, it constructs a multimodal gene representation that integrates RNA expression with two complementary data types: structural information from protein language models and functional context derived from general-purpose large language models that interpret biological knowledge. By learning a shared cell embedding space based on these macrogenes, Unify operates beyond the limitations of strict orthology. Its adversarial architecture, guided by a Pareto-optimal multi-task learning strategy, is explicitly designed to resolve the critical trade-off in comparative genomics: achieving robust batch-effect correction while simultaneously preserving the conserved biological signals essential for evolutionary discovery.

We systematically benchmarked Unify against state-of-the-art methods and demonstrated its superior ability to balance batch correction and biological conservation, even across vast evolutionary distances of over 700 million years. We showed that this integrated representation enables a range of powerful downstream applications. For instance, Unify uncovers functionally convergent gene programs that reflect critical evolutionary cellular functions, significantly improving the accuracy of predicting cellular perturbation responses between species, such as from mouse to human. By applying this framework to diverse organisms, we showed that it can construct more accurate multi-species cell-type evolutionary trees, offering a powerful new lens for comparative single-cell genomics. Overall, Unify provides a versatile and robust solution for exploring cellular function and evolutionary dynamics across the tree of life.

## Results

### Overview of Unify

To overcome the dual challenges of evolutionary divergence and technical batch effects in cross-species analysis, we developed Unify, a transfer learning framework that integrates scRNA-seq data through a novel multi-modal gene representation (**Figure 1A**). At its core, Unify moves beyond sequence homology by defining macrogenes: functionally coherent gene modules that are consistent across species. To build these, Unify first constructs a rich, multi-modal feature space for every gene by integrating two complementary sources of information: 1) structural properties, captured by embeddings from a pre-trained protein language model (ESM-2^21^), and 2) functional context, derived from a general purpose large language model (LLaMA-2^24^) interpreting curated biological knowledge. Genes with similar structural or functional embeddings are grouped into macrogenes, creating an interpretable, functional unit for integration. Unify then transfers this pre-existing, universal knowledge and applies it to the specific task of aligning cells across species.

**Figure 1.**
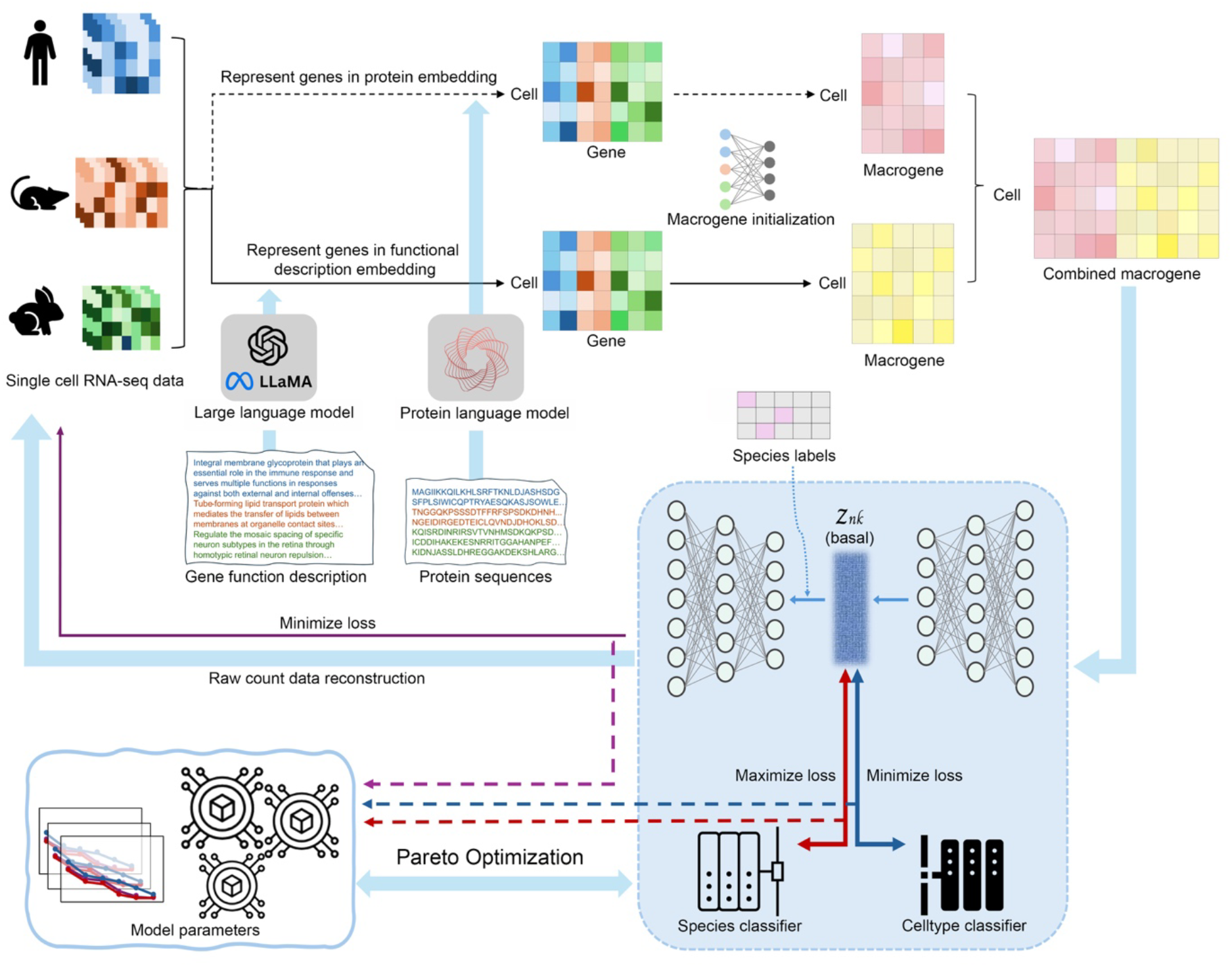
The Unify framework for balanced cross-species integration. The Unify workflow consists of two main stages. (Top Panel) First, Unify constructs multi-modal macrogenes by integrating gene representations from two complementary sources: 1) structural properties, derived from protein language models (e.g., ESM-2), and 2) functional context, derived from large language models (e.g., LLaMA-2). (Bottom Panel) These macrogenes then serve as input for an adversarial autoencoder. A Pareto-optimal multi-task learning strategy guides the training process to systematically balance three competing objectives: robust batch correction (mixing species), the preservation of conserved biological signals (aligning cell types), and the reconstruction of input data (capturing the cellular state). The final output is a unified cell embedding space that enables accurate and robust cross-species comparisons.

Before integration, Unify first projects the scRNA-seq expression profiles into a macrogene expression space. This is achieved by calculating each macrogene’s expression as a learnable, weighted combination of its constituent genes (see Methods). This resulting macrogene-by-cell matrix then serves as the input for an adversarial autoencoder, which learns the final unified latent space. This architecture is explicitly trained to disentangle biology from technology by optimizing three competing objectives simultaneously. A reconstruction loss ensures the latent space retains cellular information, while two classifiers drive the core integration: a cell-type classifier preserves conserved biological signals (bio-conservation), and a species adversarial discriminator removes technical, species-specific artifacts (batch correction).

Another key innovation in Unify is its use of Pareto-optimal multi-task learning to navigate the inherent trade-off between these objectives (**Figure 1 bottom panel**). While Pareto optimality often defines a final set of non-comparable solutions, our framework employs this principle dynamically to guide the training of a single model. Our approach, based on the MGDA-UB algorithm^25^, forgoes fixed manual weighting. Instead, at each training step, it solves for a single gradient update direction that represents the optimal compromise—a direction that improves all objectives when possible, or provides the most balanced descent when they conflict. By iteratively applying these locally optimal updates, the model follows a single, Pareto-guided training trajectory. This process yields one robust, final model, eliminating the need for any manual selection from a set of possible solutions. This principled optimization ensures that Unify achieves robust and balanced integration. The resulting framework is not only capable of generating unified cell embeddings but also allows for conditional reconstruction of gene expression, positioning it to excel at both large-scale data integration and sensitive downstream analyses. To validate its performance, we systematically benchmarked Unify against state-of-the-art methods.

### Unify Robustly Balances Batch Correction and Biological Conservation Across Vast Evolutionary Distances

To rigorously evaluate its performance, Unify was benchmarked against a comprehensive suite of competing methods. This included the current state-of-the-art in many-to-many ortholog integration, SATURN^10^ and SAMap^6^, alongside seven widely used methods^14-20^ restricted to one-to-one orthologs. Our evaluation framework was designed to assess the two fundamentals, and often competing, goals of integration: the removal of species-specific technical artifacts (batch correction) and the preservation of conserved cell-type identity (bio-conservation). Using a comprehensive suite of metrics for each aspect (Methods), we computed an overall score (40% batch correction, 60% bio-conservation) to provide a single, robust measure of integration quality, based on previous benchmark works^7, 9^.

Across all tested phylogenetic distances, from closely related species to those separated by hundreds of millions of years, Unify consistently achieved the highest overall score (**Figure 2A, Supplementary Table 1, 2**). It demonstrated a superior ability to simultaneously optimize for both batch correction and bio-conservation, a balance that other methods failed to strike (**Figure 2B-C, Supplementary Figures 1-7 for detailed integration performance in each integration task**). Crucially, while the performance of competing methods degraded significantly with increasing evolutionary distance, Unify’s performance remained remarkably stable, highlighting its robustness for challenging comparative genomics tasks.

**Figure 2.**
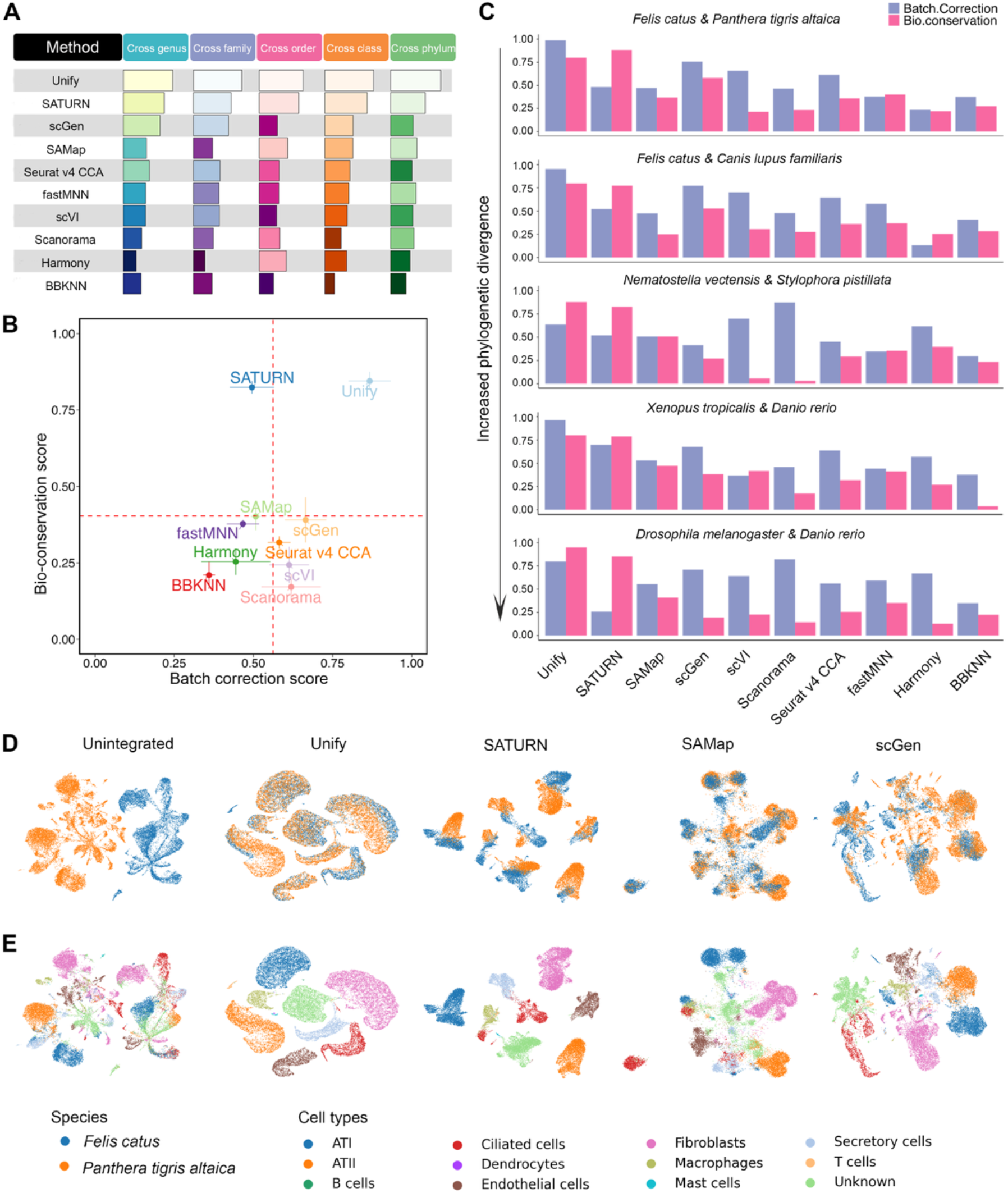
Overall performance of Unify in different cross-species integration tasks. A) Overall performance of all the benchmarked methods across different categories of cross-species integration tasks based on the phylogenetic distance between the species. B) Average bio-conservation and batch correction scores with standard error of all methods in all the integration categories. C) Integration performance of each method for different phylogenetic distance species integration tasks. D), E) UMAP plot of the selected top four methods’ integration results of integrating *Felis catus* and *Panthera tigris altaica* (cross-genus species integration task).

The specific failure modes of other methods underscore the advantages of Unify’s design. Ortholog-reliant methods like scGen predictably suffered from poor bio-conservation as the number of shared orthologs decreased significantly between distant species. Conversely, methods that incorporate all genes faced the opposite challenge. SATURN, for example, used protein embeddings to maintain strong bio-conservation but exhibited poor batch correction, failing to merge cell populations effectively at the local level (e.g., kBET, graph iLISI scores; **Figure 2D-E, Supplementary Figures 1-6**). SAMap occupied a middle ground, but its performance was inconsistent; its reliance on mutual nearest-neighbor stitching, a process sensitive to expression levels and sequencing depth^9^, resulted in suboptimal bio-conservation, particularly between closely related species.

These results offer direct support for Unify’s core architectural choices and help explain its strong performance relative to other methods. By creating multi-modal macrogenes that integrate large language model based functional annotations, Unify appears to build a more robust “functional anchor”, which likely contributes to its ability to maintain high bio-conservation. Similarly, the performance trade-offs observed across most competing methods highlight the inherent difficulty of balancing competing losses with heuristic or fixed-weight approaches. Unify’s use of Pareto-optimal adversarial training offers a more systematic approach to this problem, aiming to find a principled balance rather than a static compromise. The combination of this richer gene representation and a more principled optimization strategy likely explains Unify’s capacity to achieve a better balance between batch correction and biological discovery in cross-species integration.

### Unify enables multiple-species integration and meaningful cross-species comparison of differentially expressed macrogenes

A central goal of comparative genomics is to identify the gene programs that define cell identity and function across species. Unify moves beyond simple data integration to become an engine for this discovery through differential macrogene analysis. A key advantage of Unify is that the multi-modal macrogene approach enables the decoupling of structural and functional gene relationships, revealing evolutionary dynamics. By comparing the similarity scores from protein embeddings (structure) and text embeddings (function), we can directly identify cases of evolutionary divergence and convergence that are often missed by single-modality analysis. This allows us to systematically categorize gene pairs into groups such as: 1) conserved (high structural and functional similarity), 2) functionally divergent (high structure, low function), and 3) functionally convergent (low structure, high function). Finally, Unify overcomes the residual batch effects that confound downstream comparisons in other methods (e.g., **Supplementary Figure 8**, which shows incomplete species mixing by SATURN).

To showcase this capability, we integrated a five-species atlas of the eye’s aqueous humor outflow structures (AH Atlas) (**Figure 3B**). We then performed differential analysis at the macrogene level to identify key programs defining macrophage identity. This analysis successfully pinpointed macrogenes enriched with well-established macrophage markers along with their homologous genes (e.g., *C1QA*^*26-28*^, *C1QB*^*27, 28*^, *C1QC*^*27, 28*^, *CD74*^*29*^, *CCL3*^*28, 30*^*/4*^*28, 31*^), validating that our approach captures cross-species conserved biology (**Figure 3C)**. This finding is further substantiated by Gene Ontology (GO) and Kyoto Encyclopedia of Genes and Genomes (KEGG) analysis, which show that the component genes of these marcogenes are highly enriched in pathways characteristic of macrophage function **(Supplementary Figures 9-13)**. For instance, macrogene 857, which was highly expressed in macrophages, contained genes significantly enriched for “antigen processing and presentation,” a core macrophage function^32^ (**Figure 3D)**.

**Figure 3.**
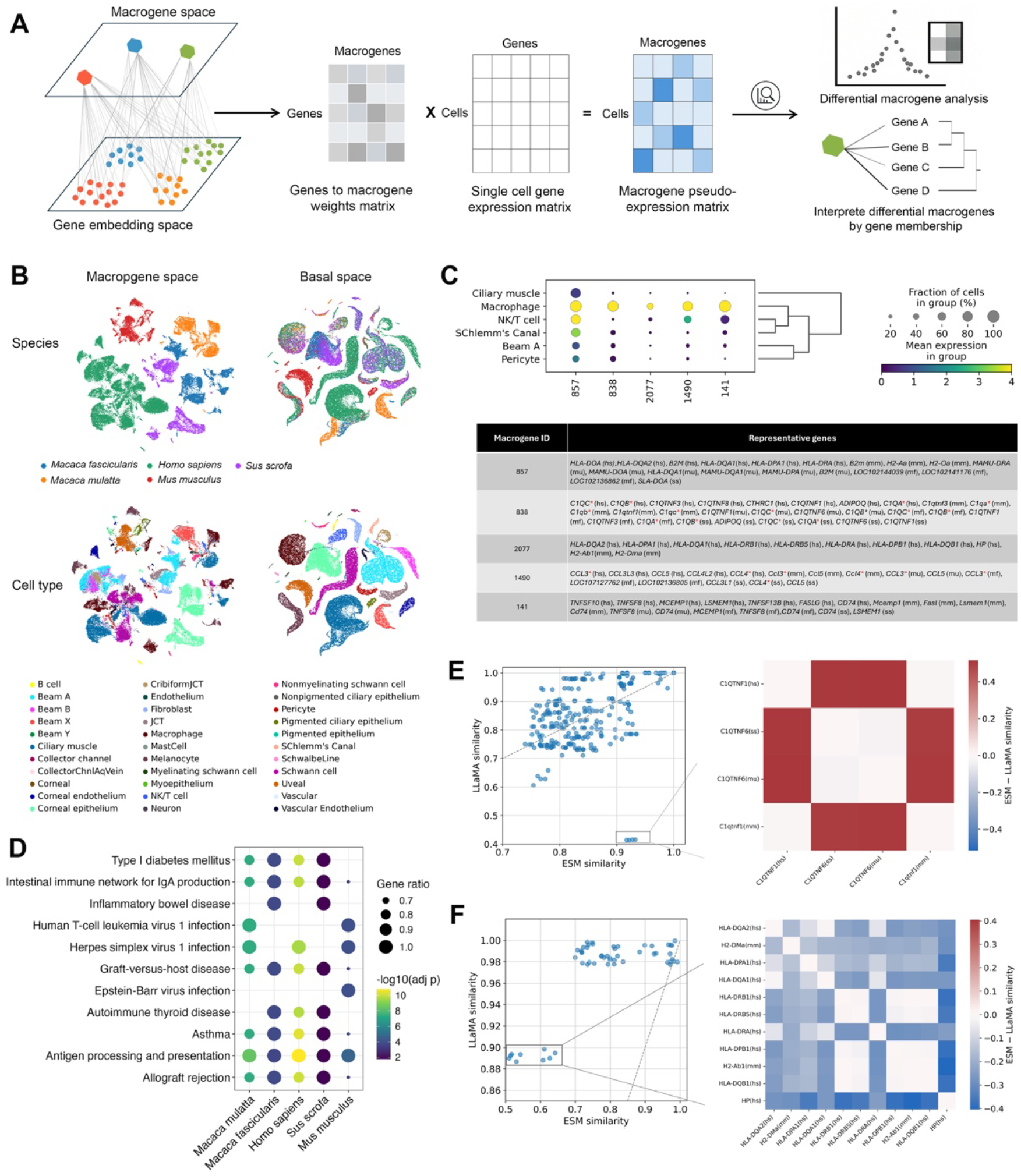
Unify integrates multi-species single-cell data to detect differential macrogene expression in the Cell Atlas of Human Trabecular Meshwork and Aqueous Outflow Structures (AH atlas), revealing genetic conservation, divergence, and convergence events. A) Schematic of Unify’s macrogene analysis. B) UMAP plot of the macrogene space and the basal space of the AH atlas. C) Dotplot of the expression levels of the differentially expressed macrogenes in macrophages (upper panel) and the component genes of each differentially expressed macrogene (lower panel). Genes marked with red stars indicate known marker genes of the macrogenes. D) The top 10 enriched KEGG pathways of the component genes of differentially expressed macrogene 857 across species in macrophage. E) The ESM and LLaMA similarity scores for gene pairs within the component genes of macrogene 838. F) Comparison of ESM and LLaMA similarity scores for gene pairs within the component genes of the differentially expressed macrogene 2077 across species of macrophages. hs: human (*Homo sapiens*); ss: pig (*Sus scrofa*); mu: monkey (*Macaque mulatta*); mm (*Mus musculus*).

A clear example of functional divergence was found in macrogene 838. This analysis revealed high structural similarity but low functional similarity between paralogous genes from the C1QTNF family, such as human C1QTNF1 and human C1QTNF6. This pattern persists across species (e.g., human C1QTNF1 vs. pig C1QTNF6), where the paralogs share a conserved protein domain but have different biological roles—one pro-inflammatory and immune-related^33^, the other anti-inflammatory and metabolic^34^ (**Figure 3E, Supplementary Figures 14-16**). This demonstrates Unify’s ability to avoid confounding functionally distinct paralogs, a critical flaw in SATURN and SAMAP, which rely on sequence similarity alone.

Conversely, we found compelling evidence of functional convergence. Macrogene 2077 contained genes like human *HP* and mouse *H2-Ab1* that share very low sequence similarity but have highly similar functional embeddings, linking them to share roles in defense and antioxidant activity (**Figure 3F, Supplementary Table 3 for the enriched GO term description of these genes**). This demonstrates Unify’s ability to identify functionally analogous genes that have evolved independently—a critical blind spot for all ortholog-based methods.

We further confirmed these findings by integrating a human and mouse immune dataset. In neutrophil, Unify successfully identified macrogenes containing neutrophil marker genes along with their orthologous genes (such as *CAMP/Camp*^*35*^, *S100A8/S100a8*^*36, 37*^, *S100A9/S100a9*^*36, 37*^, **Supplementary Figure 17 A**,**B**).

Moreover, Unify uncovered functionally coherent gene programs without direct orthologs. Macrogene 2225, for instance, contained no homologous gene pairs. However, GO enrichment analysis revealed that its component genes in both species converge on core neutrophil functions, including phosphatase activity and host-mediated cell killing^38^ (**Supplementary Figure 17C**). We further identified functionally convergent pairs, such as human *PRNP* and mouse *Calm3*. Despite sequence dissimilarity, both are implicated in calcium signaling pathways essential for neutrophil function^39^, highlighting functionally conserved programs that would otherwise be invisible, which is further illustrated by word clouds of the biological description (**Supplementary Figure 18)**.

Collectively, these results show that Unify’s differential macrogene analysis provides a powerful framework for dissecting the complex evolutionary trajectories of gene function, uncovering both conserved programs and unexpected patterns of divergence and convergence that shape cellular identity across the tree of life.

### Unify accurately predicts cross-species perturbation effects

A critical application for cross-species integration is in translational research, where predicting a human cellular response from model organism data can accelerate drug discovery and our understanding of disease. However, this task is severely hampered by methods that rely on one-to-one orthologs, which discard a vast amount of genetic information. We hypothesized that Unify’s ability to learn a unified embedding space using all genes would enable far more accurate cross-species perturbation prediction. We then evaluated the accuracy of this prediction against the actual, experimentally measured data in three keyways: 1) overall transcriptome similarity (R^2^): we calculated the R^2^ score between the predicted and actual gene expression values across all genes. A high R^2^ means the overall shape of our predicted transcriptome closely matches the real one. 2) identification of key genes (differentially expressed genes hit rate): of the genes that truly changed expression in the target species, we measured the percentage that our model correctly identified. 3) correctness of response (directional accuracy): for the genes we correctly identified, we measured how often we correctly predicted the direction of change (up-regulation vs. down-regulation).

To test this, we designed a highly challenging translational scenario: predicting the transcriptomic response of human PBMCs to IFN-β stimulation based on data from stimulated mouse lymph nodes—a setup involving different species, tissues, and experimental conditions (**Figure 4A**). When restricted to predicting changes based on one-to-one orthologs alone, Unify performed on par with state-of-the-art methods specifically developed for perturbation effect prediction, scPARM^40, 41^ (**Figure 4B**). However, the true power of Unify was revealed when the prediction was expanded beyond one-to-one orthologs to a more comprehensive gene set. By performing predictions across all highly variable genes, which capture the most dynamic aspects of the cellular response, Unify outperformed all other methods by a large margin. It achieved significantly higher accuracy (R^2^) in predicting the final expression state. It was far more successful at correctly identifying both the differentially expressed genes (DEGs) and their direction of change (**Figure 4B, C, D**). This demonstrates that Unify’s model successfully captures the complex, non-orthologous gene relationships that are essential for modeling cellular responses.

**Figure 4.**
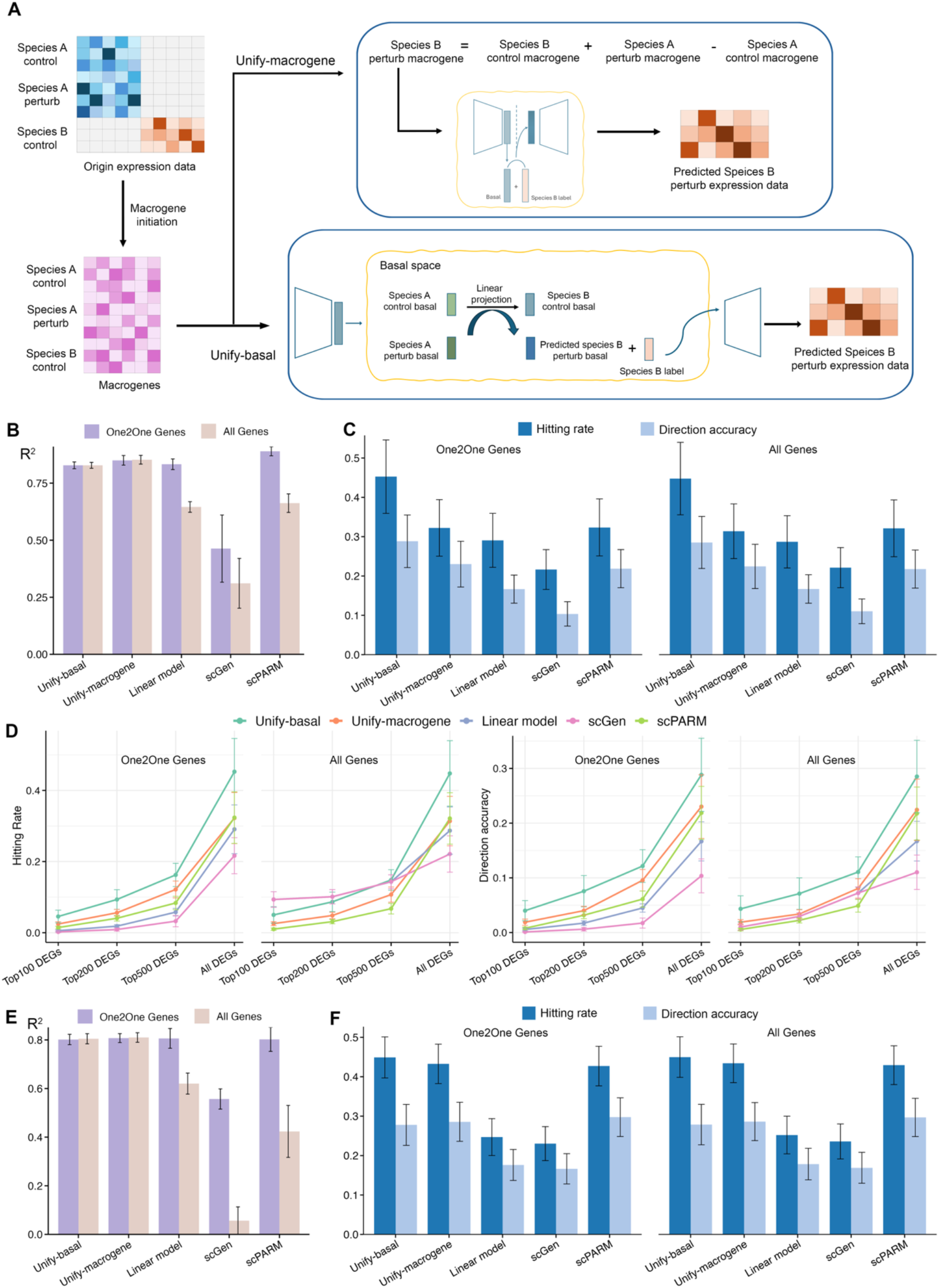
Unify accurately predicts cross-species perturbation effects. A) Schematic of the two proposed approaches for predicting perturbation effects across species using the Unify framework. B) Average R^2^ scores (with standard error) for predicted human IFN-β perturbation expression data across nine cell types. C) Average hitting rate and direction accuracy (with standard error) for differentially expressed genes predicted human in the IFN-β stimulated dataset. Left: Unify’s performance on the one-to-one orthologous gene set. Right: Unify’s performance on the all-gene set. D) Average hitting rate and direction accuracy (with standard error) of predicted differentially expressed genes by baseline methods, evaluated across different top DEGs categories in one-to-one orthologous and all-genes scenarios. E) Average R^2^ (with standard error) for predicted LPS perturbation expression data of rat, rabbit, and pig macrophages based on mouse data. F) Average hitting rate and direction accuracy (with standard error) for differentially expressed genes predicted in the LPS-stimulated rat, rabbit and pig datasets. Left: Unify’s performance on the one-to-one orthologous gene set. Right: Unify’s performance on the all-gene set.

We confirmed the robustness of this finding by performing additional cross-species predictions, using data from lipopolysaccharide (LPS) stimulated mouse macrophages to predict the responses in rat, rabbit, and pig macrophages. In every case, Unify consistently delivered superior performance, particularly when leveraging all available genes (**Figure 4E, F**). Together, these results represent a new approach for cross-species perturbation prediction. By breaking free from the constraints of one-to-one orthology, Unify provides a more comprehensive and biologically faithful framework, significantly enhancing our ability to translate findings from model organisms to human biology.

### Unify enables higher resolution cell type tree construction

One ultimate goal of cross-species integration methods is to reconstruct the evolutionary history of cell types across vast phylogenetic distances. A robust cell-type tree of life not only reveals the origins and diversification of cellular functions but also serves as a foundational step toward building comprehensive, predictive virtual cells. To demonstrate Unify’s power in this domain, we applied it to one of the most challenging integration tasks: constructing a cell-type tree of life from seven species spanning over 700 million years of evolution, including vertebrates, tunicates, insects, nematodes, and flatworms (**Figure 5A**). To evaluate the quality of this tree, we used two key metrics: 1) technical repeatability (jumble score): this is a consistency check that measures whether the tree topology is reproducible when the starting position of the maximum likelihood search is varied, ensuring it’s not a random artifact. 2) biological repeatability (scjackknife score): this is a more stringent test of robustness. It repeatedly re-builds the tree on subsets of the data to see if the core biological relationships (e.g., all vertebrate neuron types clustering together) remain stable. A high scjackknife score indicates a tree that reflects a true, reproducible biological signal^42^.

**Figure 5.**
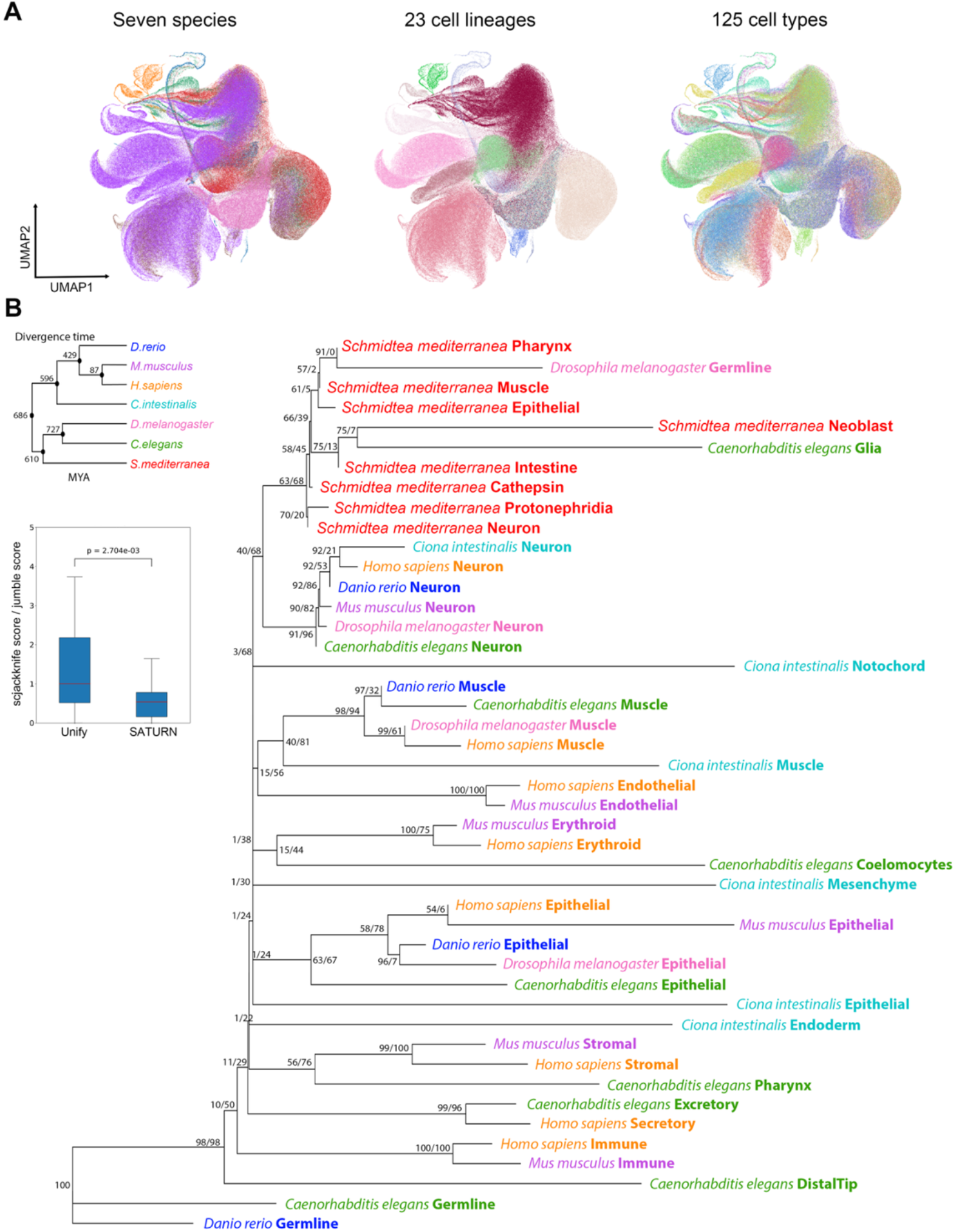
Unify enables better cell type tree construction. A) UMAP plot of the integrated single-cell datasets from seven species across the tree of life (*Schmidtea mediterranea, Danio rerio, Ciona intestinalis, Mus musculus, Homo sapiens, Drosophila melanogaster*, and *Caenorhabditis elegans*). B) Unrooted cell type tree for seven phylogenetically distant species constructed from the integrated basal embedding generated by Unify. Species and cell types are labeled at the tips. Node support values are printed as “jumble score/scjackknife score”. MYA: million years ago. Boxplot shows the distribution of the scjackknife score / jumble score for each node in the cell type tree calculated from Unify and SATURN. Two-sided Mann-Whitney U test for the biological reproducibility scores (scjackknife scores) per technical reproducibility scores (jumble scores) was conducted for the comparison.

We compared the cell-type tree generated by Unify to that of SATURN, evaluating their technical and biological repeatability. While both methods showed similar technical consistency, Unify’s tree demonstrated greater biological coherence. It achieved a significantly higher biological repeatability score, indicating that the relationships it uncovered were more consistent with known biology across independent samples (**Figure 5B, Supplementary Figure 19**). When normalized for technical performance, Unify captured substantially more meaningful biological information, demonstrating its superior ability to distill true evolutionary signals from complex, multi-species data.

By successfully organizing 125 distinct cell types from deeply divergent phyla into a biologically plausible evolutionary structure, Unify demonstrates the advantages of its design. Its multi-modal macrogenes and Pareto-optimized integration approach provide the robustness needed to bridge large evolutionary distances, moving beyond simple data alignment to deliver genuine biological insight. This high-resolution reconstruction of cell-type evolution provides a powerful new resource for understanding the fundamental principles that govern cellular identity across the animal kingdom.

## Discussion

Unify enhances cross-species single-cell analysis by tackling its two most fundamental challenges: the limitations of sequence homology and the trade-off between batch correction and biological conservation. By introducing multi-modal macrogenes that uniquely integrate structural information from protein models with functional context from large language models, Unify learns a universal cellular language that captures both functionally conserved and convergent evolution. Crucially, we pair this with a Pareto-optimal multi-task learning strategy that systematically finds the optimal balance between removing technical noise and preserving true biological signals—a principled solution to a problem that has long plagued the field.

Unify’s design unlocks previously intractable analyses by moving beyond strict one-to-one orthology. We uncovered functionally convergent gene programs that reinforce cell-type identity across species—illuminating mechanisms of evolutionary conservation in how cell types form—and simultaneously identified divergent genes whose variation underlies species-specific adaptations within the same cell type, shedding light on cellular diversification. These dual insights not only boost the accuracy of cross-species perturbation prediction—a critical step for translating model-organism findings to human disease—but also generate concrete hypotheses about the origins and evolution of cell types. A further key outcome is our 700-million-year, high-resolution cell-type tree of life, which provides a sturdy framework for tracing both convergent and divergent evolutionary trajectories. Together, these achievements establish Unify not merely as an integration tool but as a true engine of biological discovery, capable of revealing how cell types emerge, adapt, and diversify across the tree of life.

Despite its advances, Unify has limitations that open avenues for future work. Its performance is inherently tied to the quality of its underlying foundation models; as protein and text-based models become more powerful and biologically nuanced, Unify’s precision will continue to improve. While our macrogenes offer enhanced interpretability, developing automated methods for their functional annotation and characterization remains an important next step. Furthermore, extending the Unify framework to integrate other data modalities, such as epigenomic (scATAC-seq) or spatial transcriptomic data, represents an exciting frontier for building even more comprehensive cross-species atlases.

In conclusion, Unify represents a significant conceptual and technical step forward for comparative single-cell genomics. It moves the field beyond the limitations of sequence homology to a more holistic, function-centric paradigm. By providing a robust and versatile framework for integrating, interpreting, and predicting cellular behavior across the tree of life, Unify paves the way for unraveling the fundamental design principles of cell types and ultimately contributes to the ambitious goal of constructing high-resolution, predictive virtual cells^43^.

## Methods

### Overview of Unify

Unify takes multiple annotated single-cell RNA expression count datasets generated from S species *X*_*S*1_, *X*_*S*2_, …, *X*_*S*s_, where *X*_*S*i_ belongs to 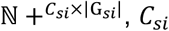 is the number of cells in species *S*_*i*_ and G_*si*_ is the set of genes in species *S*_*i*_. We used cell type annotation information sourced from the original datasets. Unify also uses two additional types of information: (1) the p-dimensional protein embeddings *P* ∈ ℝ^|*G*|×*p*^ generated from large protein language models; (2) the l-dimensional gene functional description embeddings *L* ∈ ℝ^|*G*|×*l*^ generated from the general-purpose large language models, LLaMA-2, where 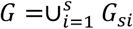.

Unify initializes *M*^*prot*^ protein-macrogenes and *M*^*func*^ functional description macrogenes by conducting the k-means clustering on the protein embedding spaces and gene functional description embedding spaces separately across all the species. Unify links the genes from multi-species to the protein-macrogenes and functional description macrogenes with weights 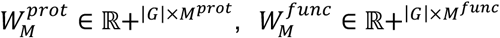, where *W*_*g,m*_, *W*_*g,n*_ ∈ ℝ +, representing the weight from a gene *g* ∈ *G* to macrogenes *m* ∈ ?^*prot*^, *n* ∈ M^*func*^. Further, for each species *S*_*i*_, Unify maps its expression data to a joint low-dimensional macrogene space by combining the protein-macrogene and functional-description macrogene together through: 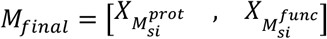 where 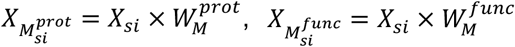.

Unify further maps the multispecies’ macrogene dataset to a joint d-dimensional latent space Z using an adversarial autoencoder framework guided by two auxiliary classifiers: a cell type classifier to promote the retention of cell type identity, while the other removes species-specific information through adversarial training. The resulting latent embedding captures conserved cellular features across species. We further applied conditional decoding based on given species labels to accurately reconstruct either the original gene expression profiles or the macrogene dataset. To balance competing objectives: preserving biological signals, removing technical batch effects and ensuring faithful reconstruction, we used the MGDA-UB multi-task learning algorithm^25^, aiming to identify model parameters that achieve an optimal trade-off across these goals.

### Generation of protein embeddings using a protein language model

Protein embeddings were generated using a pretrained protein embedding language model applied to the reference proteome of each species (Supplementary Table 4 for the proteome information used for each species in this study). The protein embedding language model takes the protein’s amino acid sequence as input and outputs a dimensional vector representing the embedding of the protein. For benchmarking, protein embeddings were generated using the ESM-1b^44^ model; in all other analyses, we employed the updated ESM-2^21^ model for embedding generation. Specifically, the ESM-1b model outputs a dimensional vector of 1280 to represent the protein, while ESM-2 model outputs a dimensional vector of 5120 to represent the protein. The gene embeddings used in the study are the average of all the possible proteins source from the same gene. Any other protein language models that can reflect the sequence characteristics of the coding gene can be used as an input to Unify.

### Generation of gene function description embeddings from general-purpose large language model

Gene functional description embeddings were generated from the general-purpose pretrained large language model using each species’ Gene Ontology functional description for each gene sequence. The large language model takes the text description as input and outputs a dimensional vector representation of the embedding of the genes’ function. We used the open-source LLaMA-2^24^ model to generate the functional description embedding of each gene in the species. The gene functional description can either be downloaded from the NCBI website^45^, Gene Ontology website^46^, Uniprot database^47^ or generated de novo based on the protein sequences (Supplementary Note 1). Apart from the Gene Ontology description, any other term that describes the gene function, such as KEGG pathways, Reactome pathways, could be used to generate the gene functional description embeddings. We envision that any other general-purpose language model could be used to generate embeddings that work as inputs to Unify.

### Macrogene initialization

To capture gene-level similarity from both structural and functional perspectives, we constructed two sets of macrogenes: one based on protein embeddings and the other based on functional description embeddings. These macrogenes serve as interpretable meta-units representing groups of genes with shared sequence or functional properties.

### Protein-based macrogene construction

For each gene *g*_*si*_ from species *S*_*i*_, we obtained its average protein embeddings 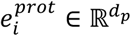, derived from pretrained protein language models. All gene embeddings across species are concatenated into a joint matrix:

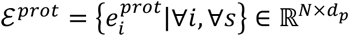

We applied K_p_-means clustering to this matrix to identify K_p_ protein-based macrogenes, where each cluster centroids 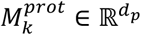.

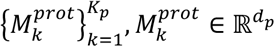

### Functional-based macrogene construction

Similarly, we generated functional description embeddings 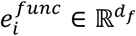 using the pretrained large language model based on gene function descriptions. They are aggregated as:

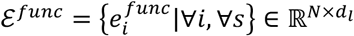

We performed K_f_-means clustering on ℰ^*func*^ to generate K_f_ functional macrogenes, where each cluster centroid 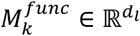.

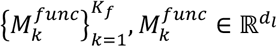

### Gene-to-macrogene weight initialization

For both protein and functional macrogenes, we computed a soft assignment weight matrix *W* ∈ ℝ^*N*×*K*^, where *W*_*gk*_ represents the contribution of gene g to macrogene k. These weights are initialized using a ranked Euclidean distance between each gene and all the macrogene centroids. Let *rd*_*g,k*_ ∈ {1,2, …, R} denotes the rank of all the genes g for a specific macrogene k. *rd*_*g,k*_ = 1 indicates the nearest gene to macrogene k. The initial weight is defined as:

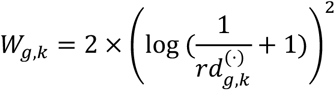

where (·) ∈ {*prot, func*}.

### Macrogene expression initialization

Given the raw gene expression matrix 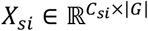 for species *S*_*i*_, where *C*_*si*_ is the number of cells, and |*G*| is the number of genes, the macrogene expression matrix 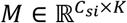 is computed as:

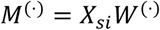

for protein-based and functional-based macrogenes, respectively.

### Combined macrogene representation

The two macrogenes’ expression matrices are concatenated to form the combined macrogenes expression representation:

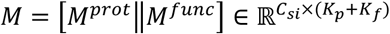

This combined representation serves as input to the downstream component of Unify.

### Adversarial learning and Pareto optimization during training with an autoencoder

After obtaining the combined macrogene representation for each species, Unify employs an adversarial autoencoder to learn species-invariant but biologically informative embeddings of single cell transcriptomes. The training process is designed to ensure that the latent representation retains meaningful cell-type information while discarding species-specific variation. This is achieved through a combination of three loss terms: reconstruction loss, cell-type classification loss and adversarial species classification loss, which are balanced through Pareto optimization.

### Autoencoder Framework

Given a transcriptome expression profile X_c_ for a cell c, we computed a macrogene activation vector e_c_ by applying a transformation based on gene-to-macrogene weights:

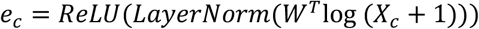

Here, *W* ∈ ℝ^|*G*|×|*M*|^ is the gene-to-macrogene weight matrix, and LayerNorm refers to standard layer normalization:

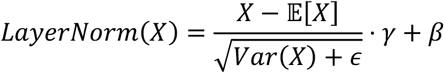

The macrogene representation *e*_*c*_ ∈ ℝ^|*M*|^ is then passed through a four-layer neural network encoder with ReLU activations, layer normalizations and dropout to produce a low-dimensional basal embedding:

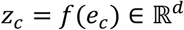

Further, the decoder De reconstructs either the original macrogene representation or the expression profile from the basal embedding, conditioned on the species label s_c_:

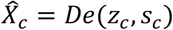

The reconstruction loss is defined as:

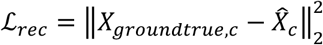

### Adversarial training

To enforce biological relevance and species-invariance in the embedding space, we introduced two auxiliary classifiers:

1. A cell type classifier that encourages cell identity by minimizing the cell-type classification loss:

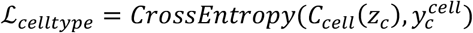
2. A species classifier trained adversarially to remove species-specific signals by maximizing species prediction error:

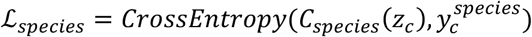

The overall loss during training is expressed in the equation below:

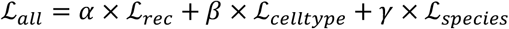

A naive approach could be optimizing a fixed set of alpha, beta and gamma. However, manually tuning these weights is subjective and fails to capture the dynamic trade-offs during training.

To resolve this, we framed the problem as multi-objective optimization and employed the Multiple Gradient Descent Algorithm - Upper Bound (MGDA-UB)^25^ to find a Pareto-optimal solution. Instead of using static weights, MGDA-UB treats each loss as a distinct objective. At each training step, it computes the gradients of each loss with respect to the shared latent cell embeddings. It then solves for a common descent direction that represents the best trade-off among the competing objectives, effectively finding weights that steer the model towards the Pareto front—the set of solutions where one objective cannot be improved without degrading another. This principled approach removes the need for arbitrary loss weighting and ensures that Unify robustly and automatically balances the competing goals of bio-conservation and batch correction while retaining its ability to reconstruct the input data.

### Perturbation prediction using Unify

Unify supports two strategies (**Unify-Macrogene** and **Unify-basal approach**) for predicting perturbation responses from a source species to a target species.

### Unify-Macrogene Approach

Unify-macrogene is an approach based on macrogene spaces, which encodes abstracted gene modules from both structural and functional features learned from both species. We used the below function to predict the target species’ macrogene under perturbation:

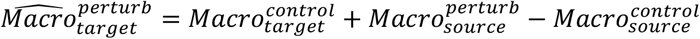

where 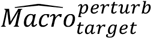 is the predicted macrogenes of the target species under perturbation; 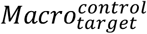 is the macrogenes of the target species under control condition. 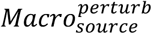 is the macrogenes of the source species under perturbation. 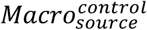 is the macrogenes of the source species under control condition.

Once 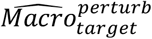 is obtained, it is then passed through Unify’s decoder to generate the corresponding perturbed gene expression profile of the target species.

### Unify-Basal Approach

In this approach, a linear transformation F is trained to align the basal embeddings of the control condition from the source species to the target species.

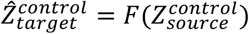

After the model was trained, we used the model to predict the basal embedding under the perturbed condition of the target species from the source species.

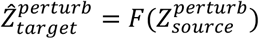

The transformed embeddings 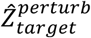 are directly input into Unify’s decoder to obtain the predicted perturbation-response expression profile in the target species.

Both strategies leverage Unify’s adversarially learned, species-invariant decoder, yielding accurate, high-resolution predictions of how perturbations translate across evolutionary distances.

### Statistics

We used SCANPY^48^ to perform the Wilcoxon rank-sum test for calculating the differentially expressed macrogenes cross species. Specifically, we used the *rank_genes_groups* function with default model parameters.

The R^2^ between the ground truth gene expression profile and the predicted one is calculated using the function *r2_score* from sklearn.metrics^49^.

Two-sided Mann-Whitney U test is conducted for comparing the biological reproducibility scores (scjackknife scores) per technical reproducibility scores (jumble scores) extracted from the unrooted cell type tree for seven phylogenetically distant species of SATURN and Unify. We used the function *mannwhitneyu* from scipy.stats^50^.

### Determining gene homologs

Gene ortholog information was retrieved using the Ensembl BioMart tool^51^ via the Ensembl genome browser^52^.

### Benchmark evaluations

#### Methods description

We evaluated Unify, SATURN^10^, SAMap^6^, scGen^17^, scVI^18^, Harmony^19^, fastMNN^15^, BBKNN^20^, Scanorama^16^ and Seurat v4 CCA^14^ in the different phylogenetic distance species integration tasks. Among these methods, Unify, SATRUN and SAMap accept all genes from the integrated species as input, while the other methods only take the one-to-one orthologous genes between the integrated species as input. For more details about each selected method, see Supplementary Note 2.

#### Evaluation metrics

For cross-species datasets integration purposes, we evaluate the methods’ performance in batch-correction and bio-conservation scores using scIB packages^7^. Batch-correction scores are calculated based on the average of below sub-metrics: batch ARI, batch ASW, graph iLISI, batch NMI, kBET, PCR batch and graph connectivity. Bio-conservation scores are calculated based on the average of the sub-metrics: cell type ARI, cell type ASW, graph cLISI, cell type NMI, HVG conservation and trajectory conservation. We further combined batch-correction scores with bio-conservation scores in a ratio of 4:6 to form an overall score for direct comparison (Supplementary Note 3).

### Protein sequences comparison

To analyze and compare gene sequence conservation, we employed a multi-step approach. First, protein sequences of the target genes were aligned using CLUSTALW^53^ to perform multiple sequence alignment (MSA), enabling the identification of conserved and variable regions across sequences. The aligned sequences were then input into the SWISS-MODEL^54^ homology modeling server to predict the three-dimensional (3D) protein structures, which were returned in PDB format. Finally, we used ESPript^55^ (Easy Sequencing in PostScript) to integrate the MSA results with the predicted 3D structures. This tool enabled the visualization of secondary structural elements (e.g., α-helices, β-strands) and facilitated a detailed comparison of both sequence and structural differences among the selected proteins.

### GO term and KEGG enrichment analysis

To investigate the biological functions and pathways associated with the component genes of the differentially expressed macrogenes, we performed Gene Ontology (GO) term and Kyoto Encyclopedia of Genes and Genomes (KEGG) pathway enrichment analyses. Differentially expressed macrogenes were identified based on predefined threshold-adjusted p-values less than 0.05. For GO enrichment, we focused on the Biological Process (BP) category to capture functional aspects of gene regulation.

Enrichment analysis was conducted using the clusterProfiler^56^ R package (version 4.10.1). Background gene sets were defined based on all expressed genes in the corresponding dataset. The enrichment results were calculated using a hypergeometric test, and multiple testing correction was applied using the Benjamini–Hochberg method. Terms with adjusted p-values (false discovery rate, FDR) below 0.05 were considered significantly enriched.

For KEGG analysis, we used gene identifiers mapped to KEGG orthology via the org.X.eg.db annotation package (where “X” corresponds to the organism, specifically, *Hs* for human or *Mm* for mouse, Ss for pig, Mmu for *Macaque mulatta*, Mmf for *Macaca fascicularis*). KEGG pathway enrichment was performed using the same statistical framework as GO analysis. The enriched GO terms and KEGG pathways were visualized using dot plots for interpretability.

### Cell type tree construction

To construct the cell type phylogenetic tree spanning seven species (*Schmidtea mediterranea, Danio rerio, Ciona intestinalis, Mus musculus, Homo sapiens, Drosophila melanogaster*, and *Caenorhabditis elegans*), we followed the approach described by Jasmine et al^42^. First, we generated integrated cross-species embeddings. Cell types represented by fewer than 100 cells were excluded to ensure robustness. We then performed principal component analysis (PCA) on filtered data and retained the top 20 principal components for downstream analysis. To reduce computational complexity, one cell per cell type per species was randomly sampled. We constructed the phylogenetic tree using the contml program from the PHYLIP package (version 3.698), applying a Brownian motion model. Finally, to evaluate both technical and biological reproducibility of the resulting tree, we calculated jumble scores and scjackknife scores for each tip by randomly repeating the tree construction process and randomly selecting the starting tip of the tree for 500 times separately (Supplementary Note 4).

## Data and code availability

Datasets are collected from published studies, as described in Supplementary Table 1. The output results for the benchmark session and cross-species perturbation prediction session are available from Supplementary Table 2 and Supplementary Table 5-6, respectively. The source code is available at https://github.com/huawen-poppy/Unify.git.

