## Supplementary_Figures for "Unify: Learning Cellular Evolution with Universal Multimodal Embeddings"

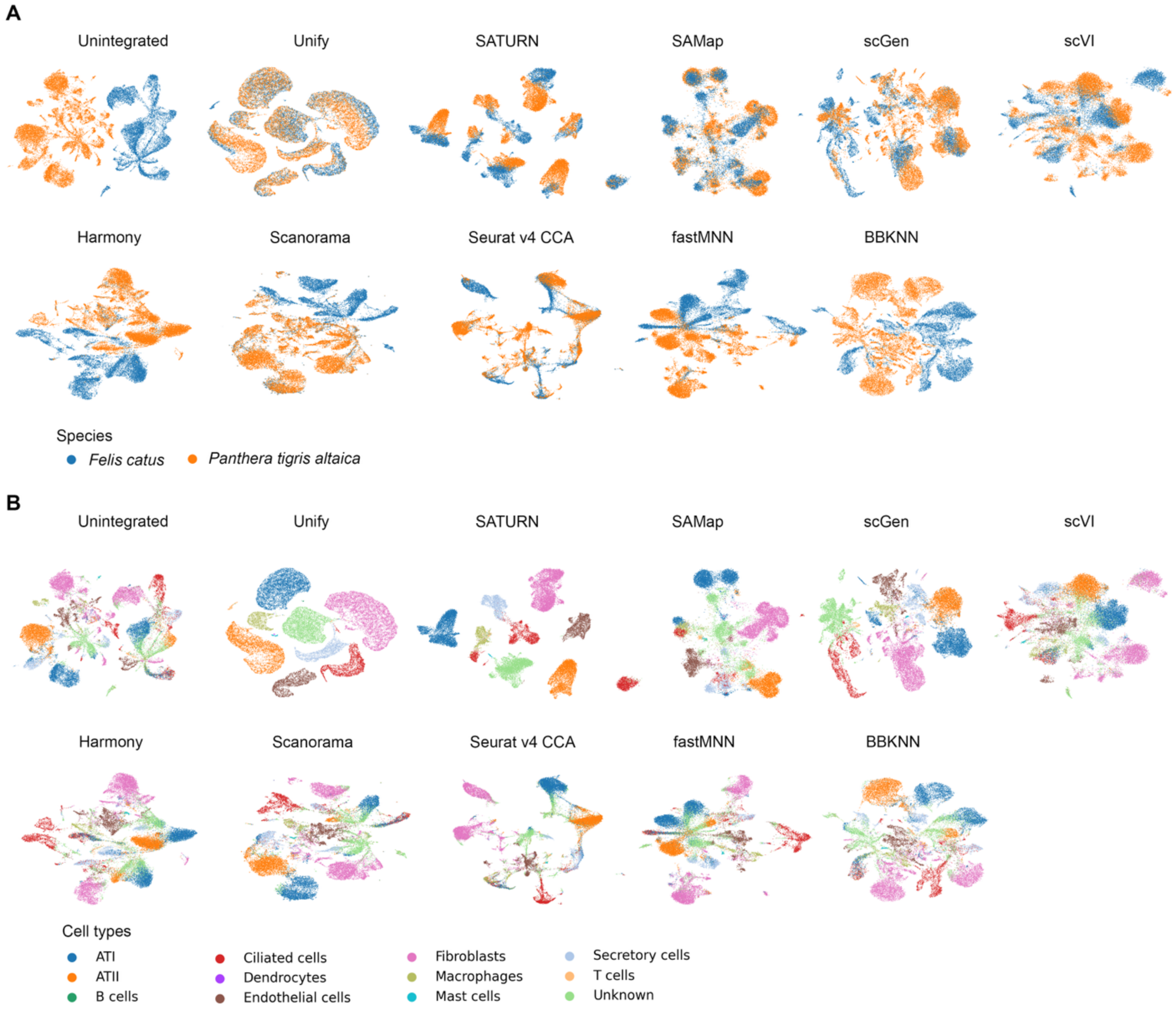


**Figure S1.** UMAP layouts for the unintegrated and integrated *Felis catus* (cat) and *Panthera tigris altaica* (tiger) datasets from different tissues colored by species labels (**A**) and colored by cell type labels (**B**).

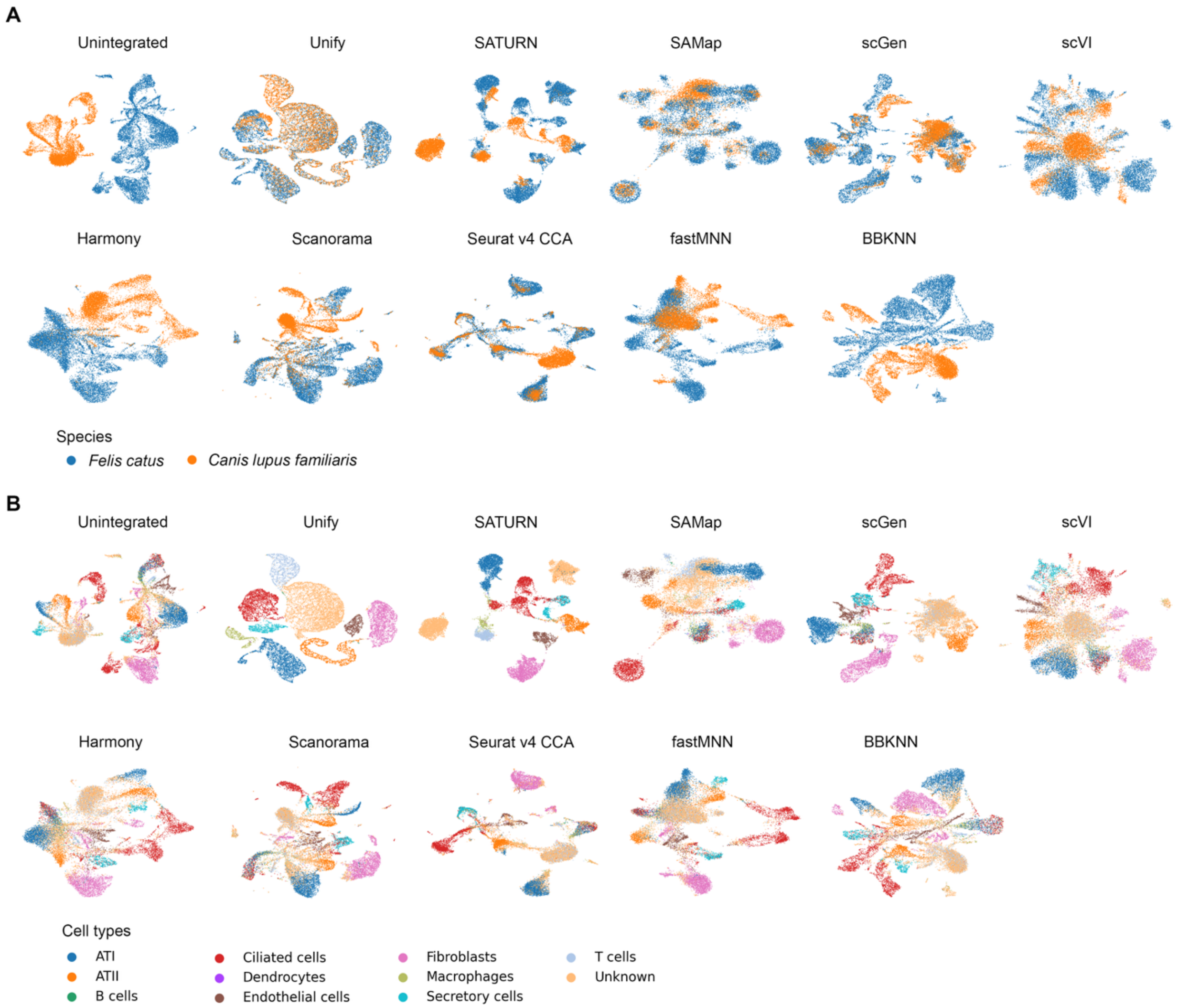


**Figure S2.** UMAP layouts for the unintegrated and integrated *Felis catus* (cat) and *Canis lupus familiaris* (dog) datasets from different tissues colored by species labels (**A**) and colored by cell type labels (**B**).

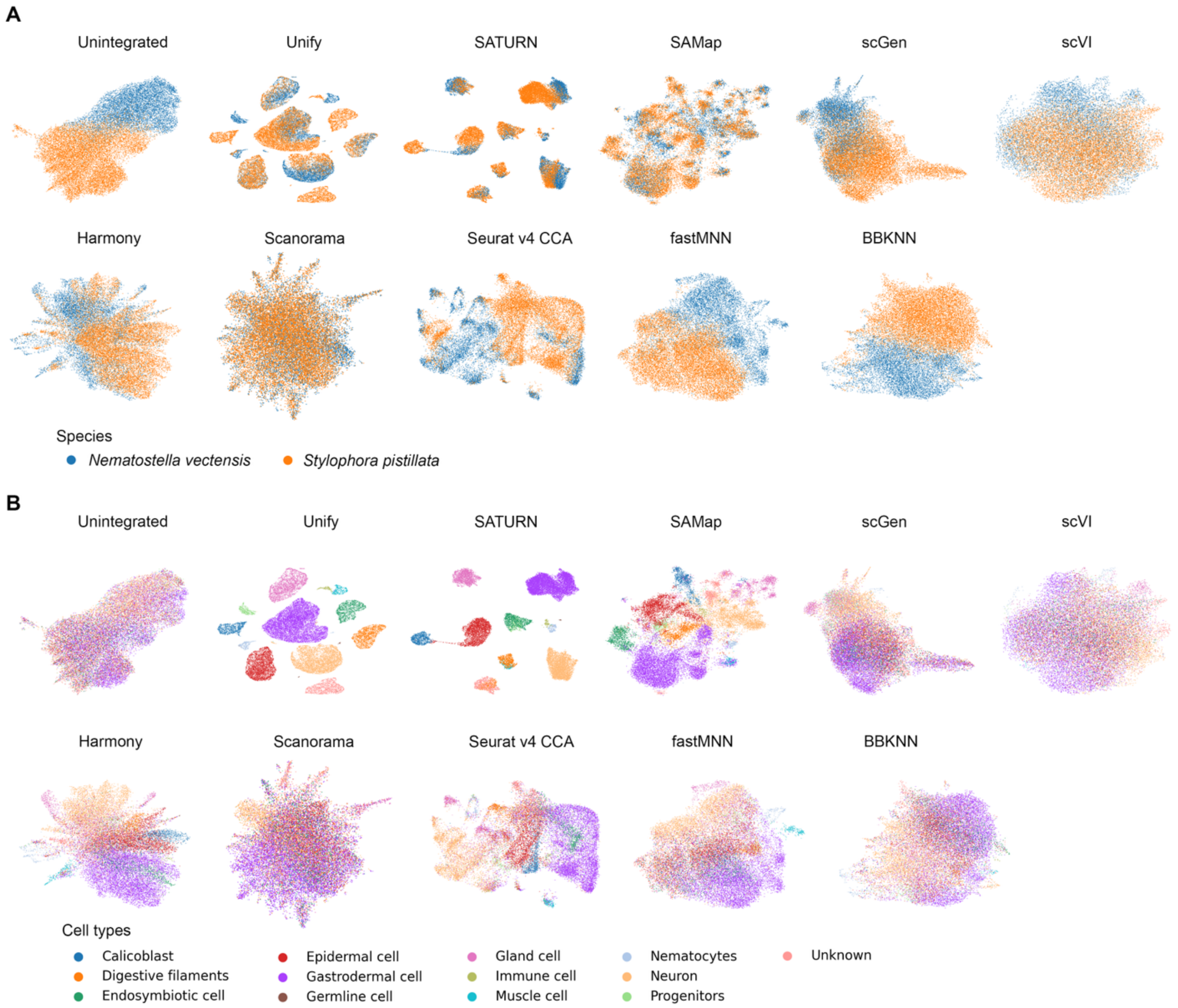


**Figure S3.** UMAP layouts for the unintegrated and integrated *Nematostella vectensis* (sea anemone) and *Stylophora pistillata* (hard coral) datasets. (**A**) Plots are colored by species labels. (**B**) Plots are colored by cell type labels.

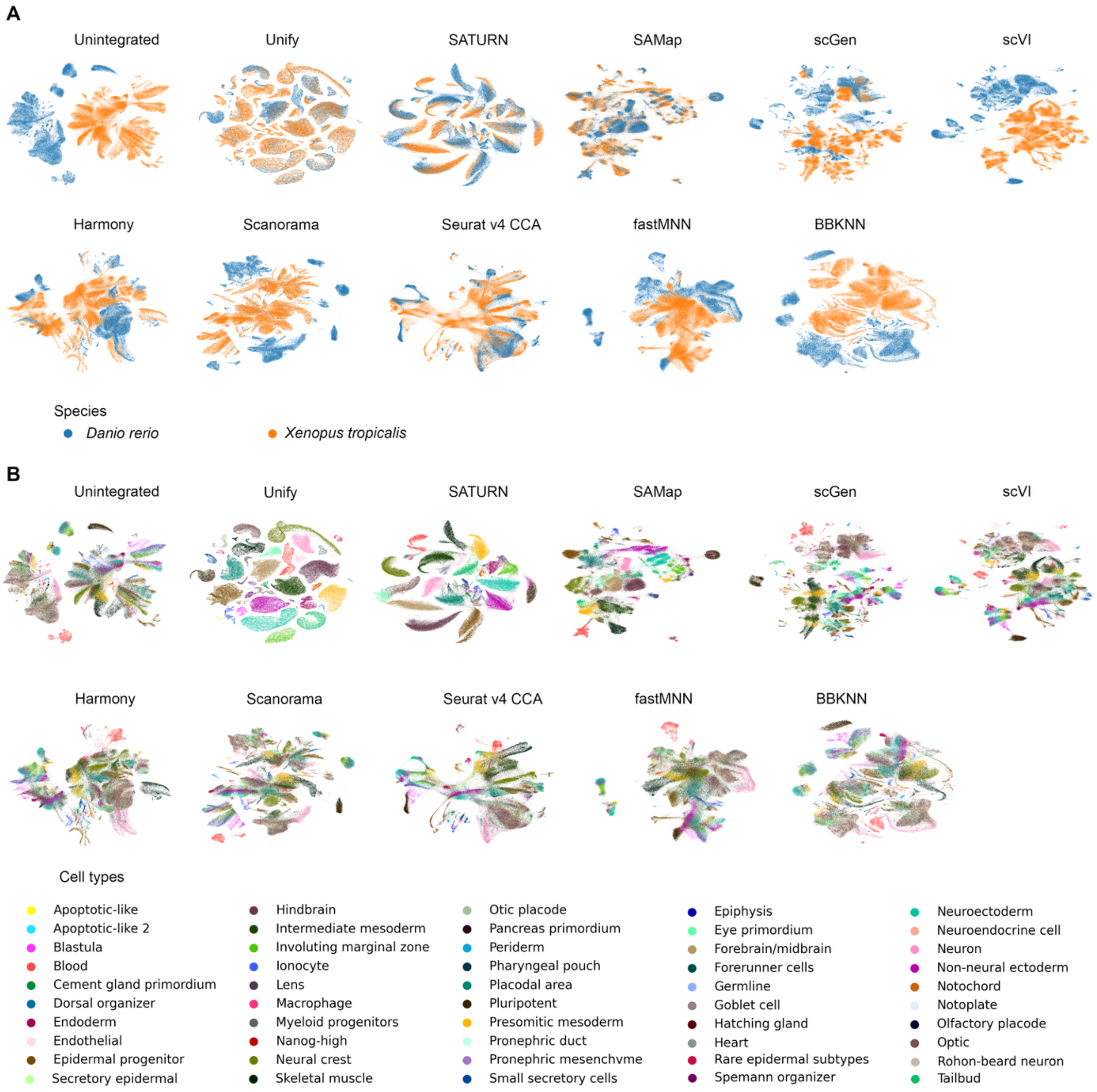


**Figure S4.** UMAP layouts for the unintegrated and integrated *Danio rerio* (zebrafish) and *Xenopus tropicalis* (frog) datasets. (**A**) Plots are colored by species labels. (**B**) Plots are colored by cell type labels.

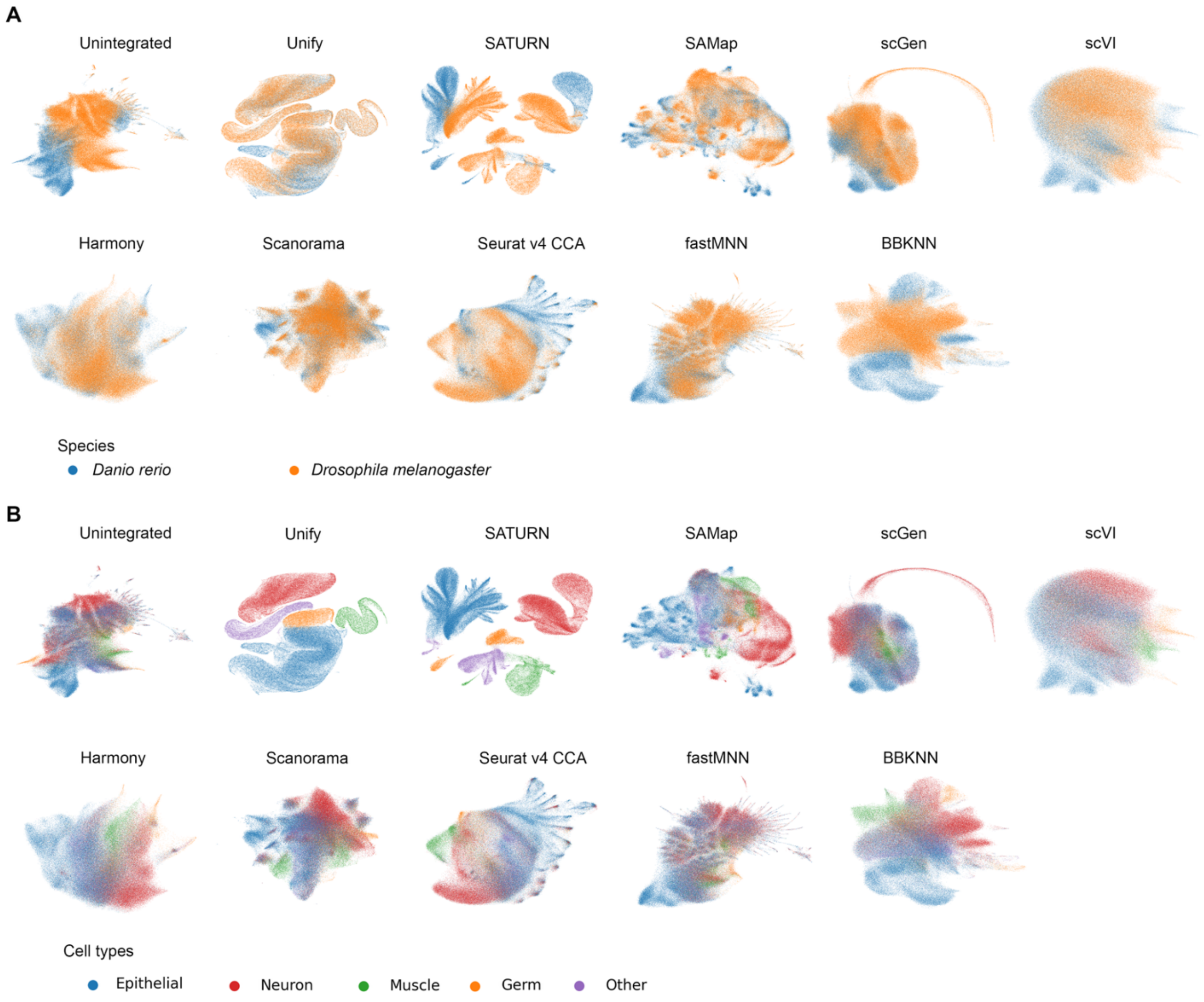


**Figure S5.** UMAP layouts for the unintegrated and integrated *Danio rerio* (zebrafish) and *Drosophila melanogaster* (fly) datasets. (**A**) Plots are colored by species labels. (**B**) Plots are colored by cell type labels.

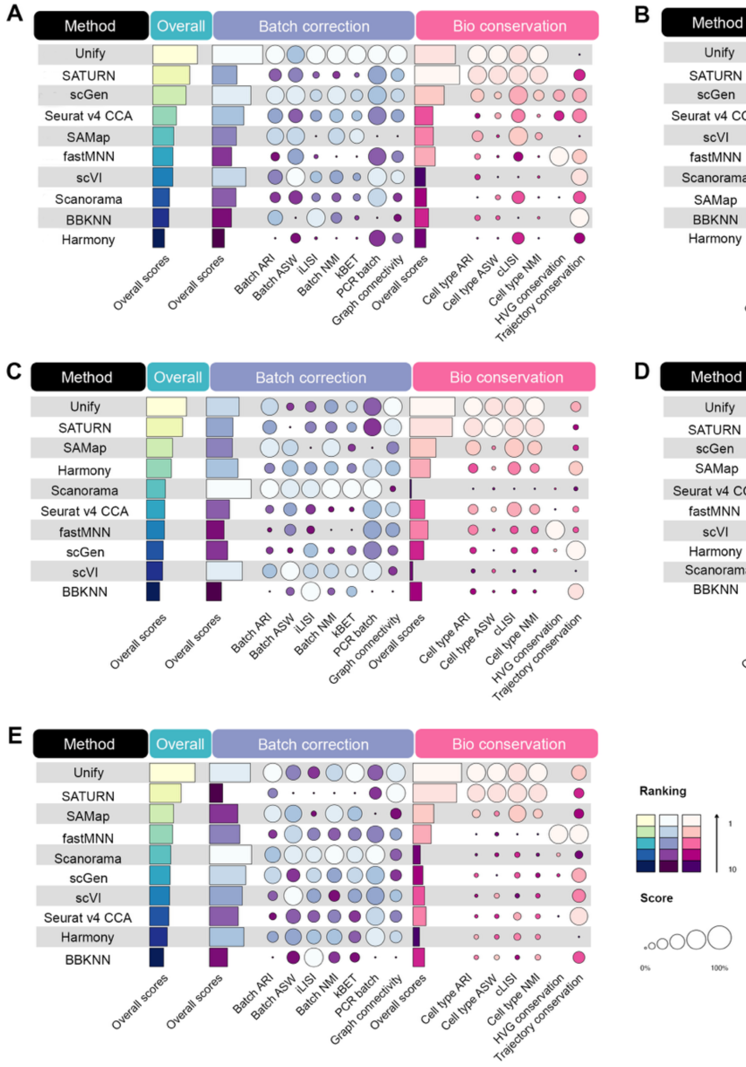


**Figure S6.** Overview of the benchmarking results for different species integration categories. (**A**) Ranking result for the cross-genus species (*Felis catus* (cat) and *Panthera tigris altaica* (tiger)) integration result. (**B**) Ranking result for the cross-family species (*Felis catus* (cat) and *Canis lupus familiaris* (dog)) integration result. (**C**) Ranking result for the cross-order species (*Nematostella vectensis* (sea anemone) and *Stylophora pistillata* (hard coral)) integration result. (**D**) Ranking result for the cross-class species (*Danio rerio* (zebrafish) and *Xenopus tropicalis* (frog)) integration result. (**E**) Ranking result for the cross-phylum species (*Danio rerio* (zebrafish) and *Drosophila melanogaster* (fly)) integration result.

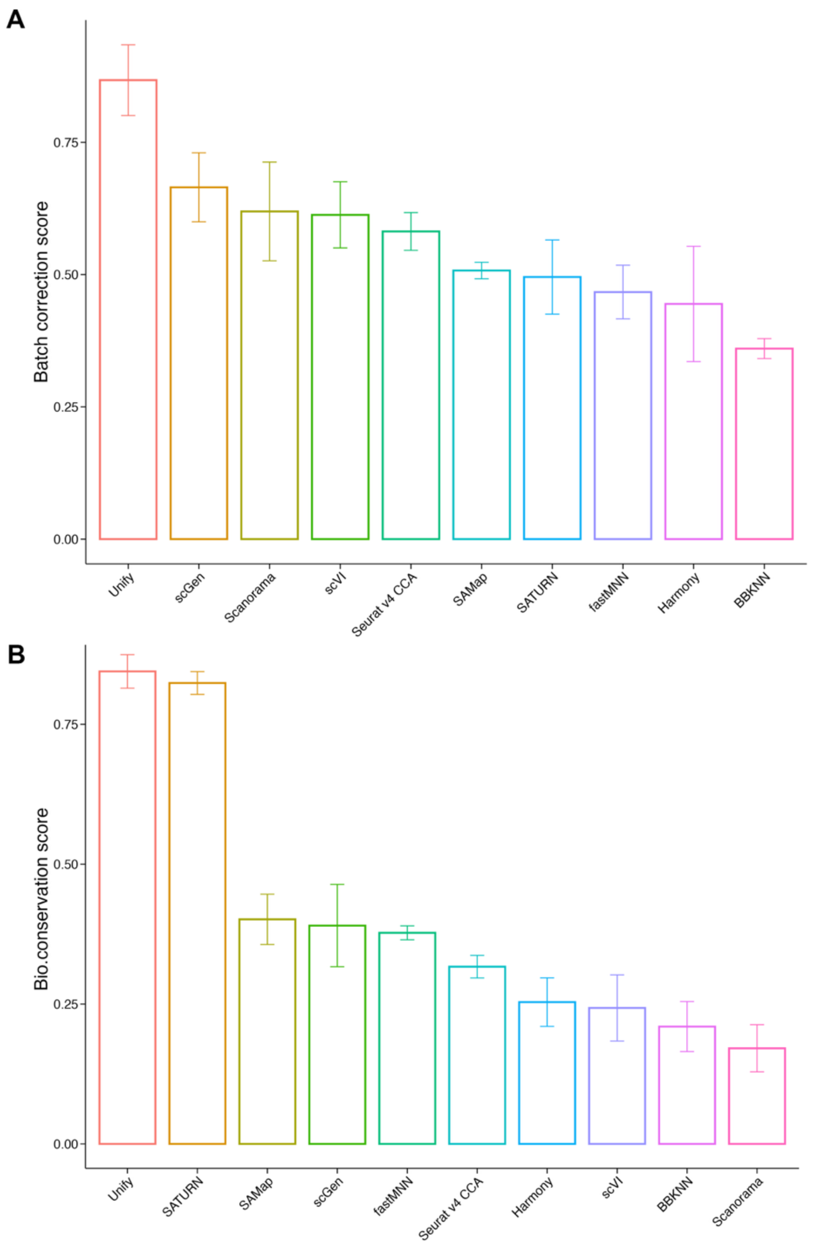


**Figure S7.** Overall batch correction scores and bio-conservation scores for cross species integration tasks. (**A**) Bar plot of the overall batch correction scores for ten methods in all tasks. (**B**) Bar plot of the overall bio-conservation scores for ten methods in all tasks.

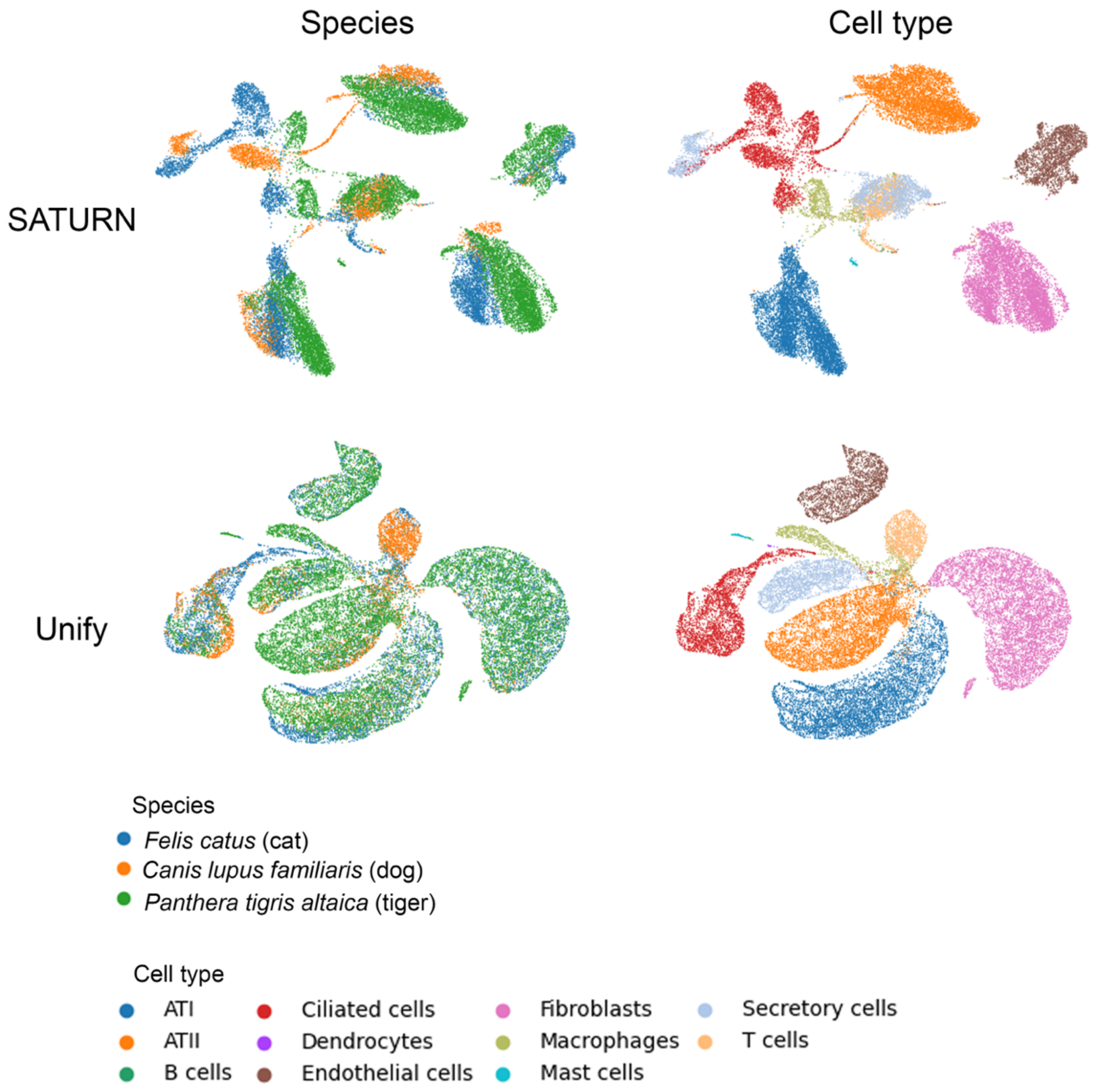


**Figure S8.** UMAP layouts for the results from SATURN and Unify of integrating *Felis catus* (cat), *Canis lupus familiaris* (dog) and *Panthera tigris altaica* (tiger) datasets. SATURN fails to mix the species well for some cell types, such as Ciliated cells and Secretory cells.

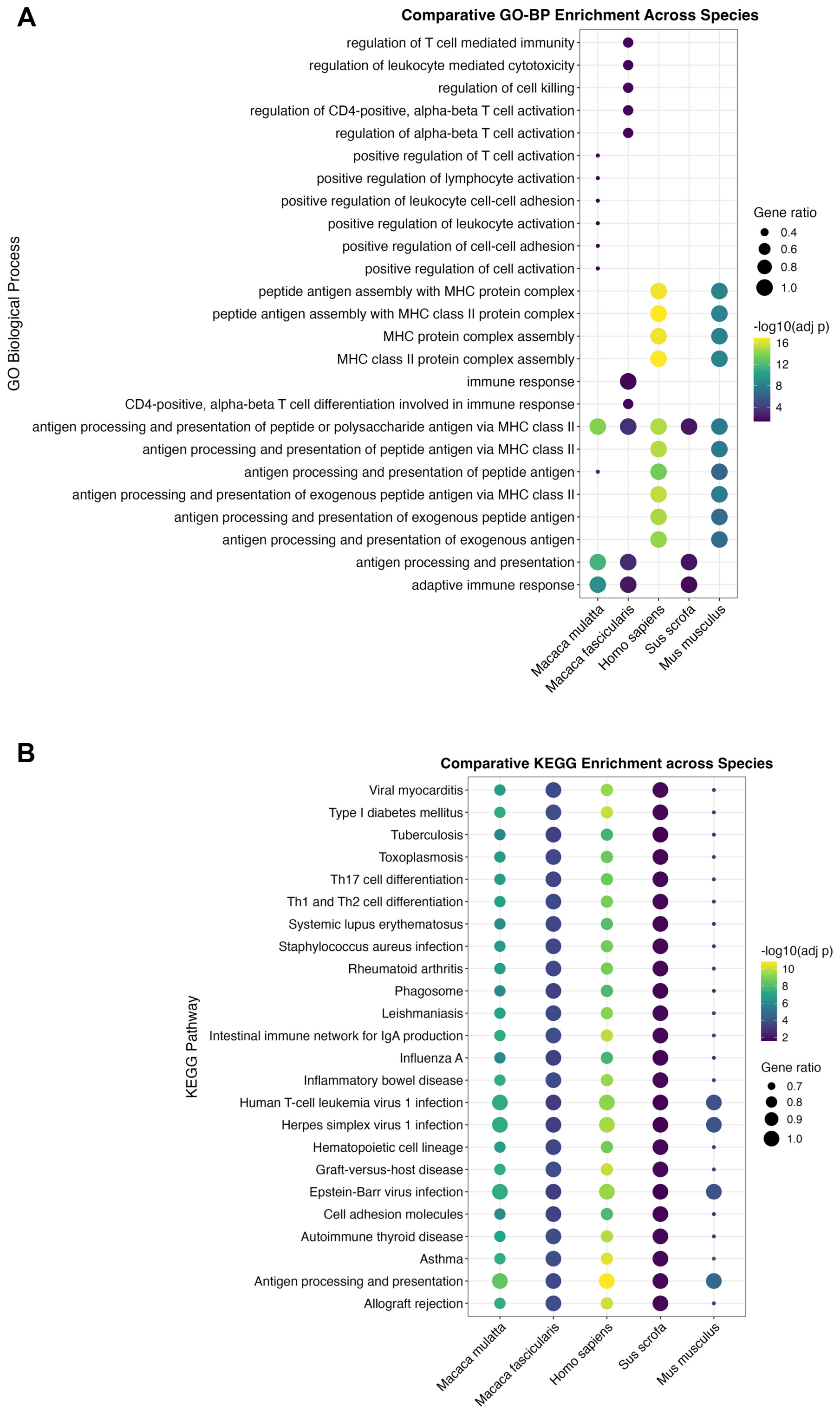


**Figure S9.** Gene Ontology term enrichment and KEGG enrichment plots for the component genes of macrogene 857 for *Macaca mulatta, Macaca fascicularis, Homo sapiens, Sus scrofa* and *Mus musculus* in the AH atlas. (**A**) Gene ontology term enrichment plot. (**B**) KEGG enrichment plot.

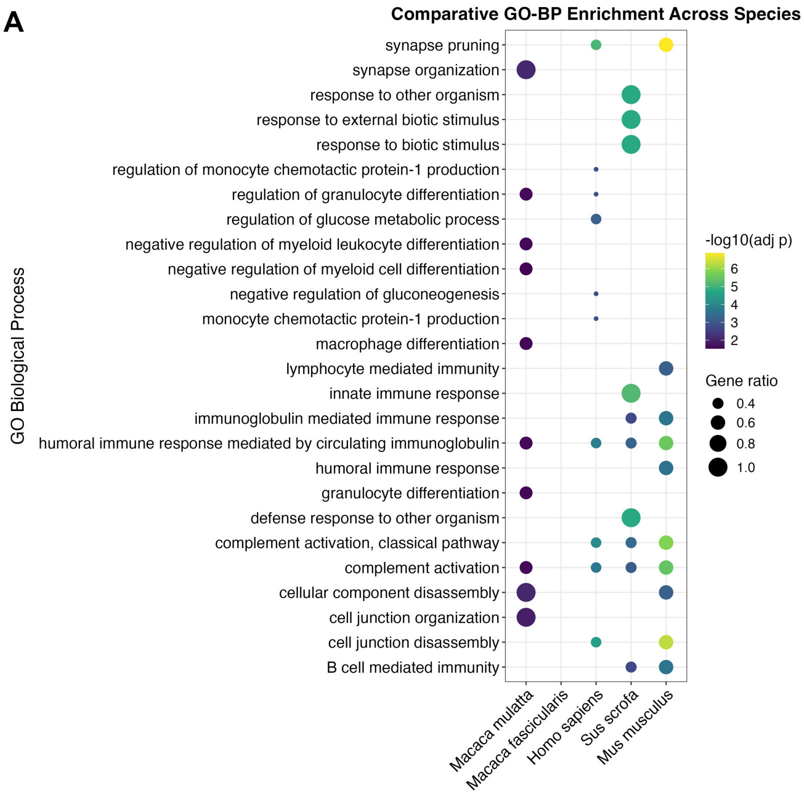


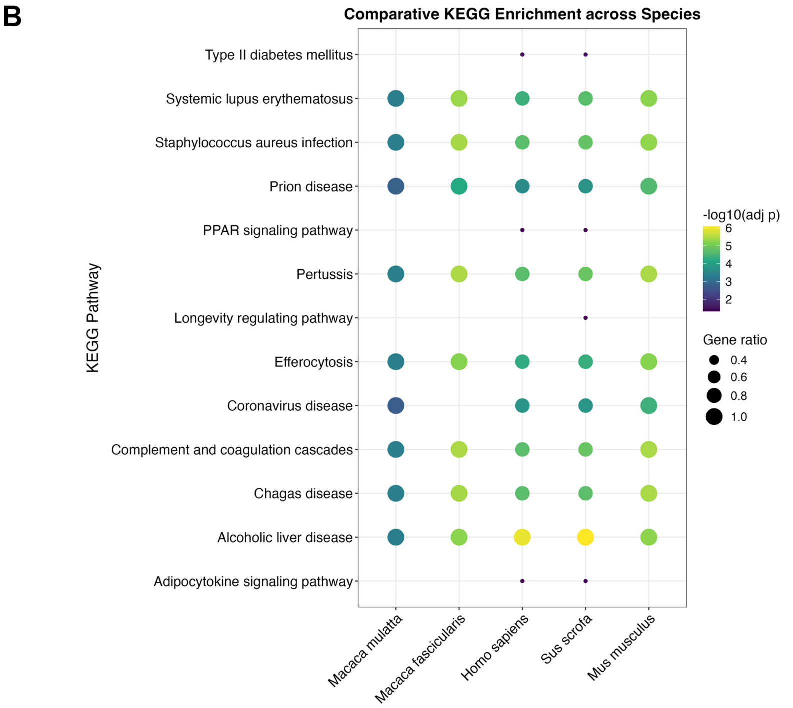


**Figure S10.** Gene Ontology term enrichment and KEGG enrichment plots for the component genes of macrogene 838 for *Macaca mulatta, Macaca fascicularis, Homo sapiens, Sus scrofa* and *Mus musculus* in the AH atlas. (**A**) Gene ontology term enrichment plot. (**B**) KEGG enrichment plot.

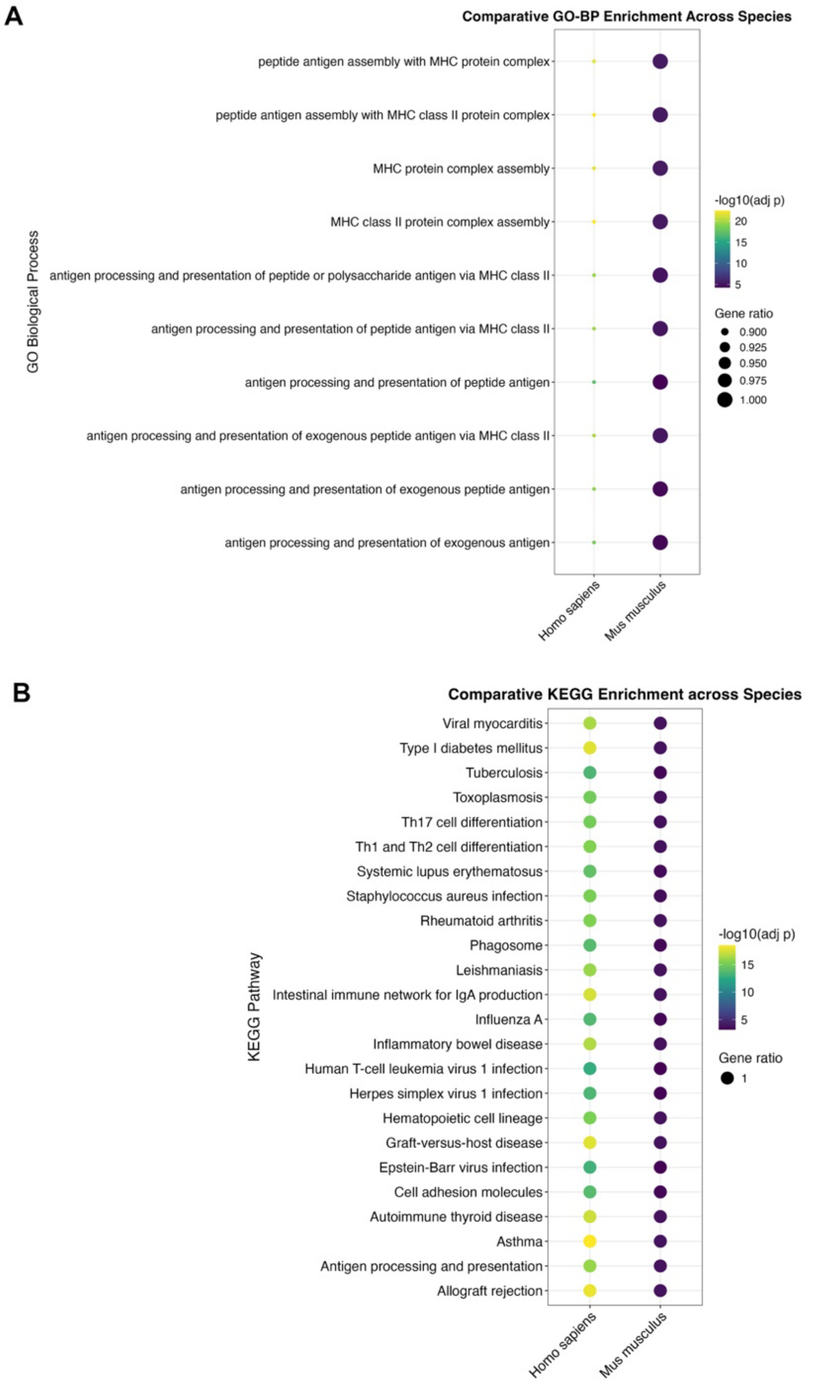


**Figure S11.** Gene Ontology term enrichment and KEGG enrichment plots for the component genes of macrogene 2077 for *Macaca mulatta, Macaca fascicularis, Homo sapiens, Sus scrofa* and *Mus musculus* in the AH atlas. (**A**) Gene ontology term enrichment plot. (**B**) KEGG enrichment plot.

A

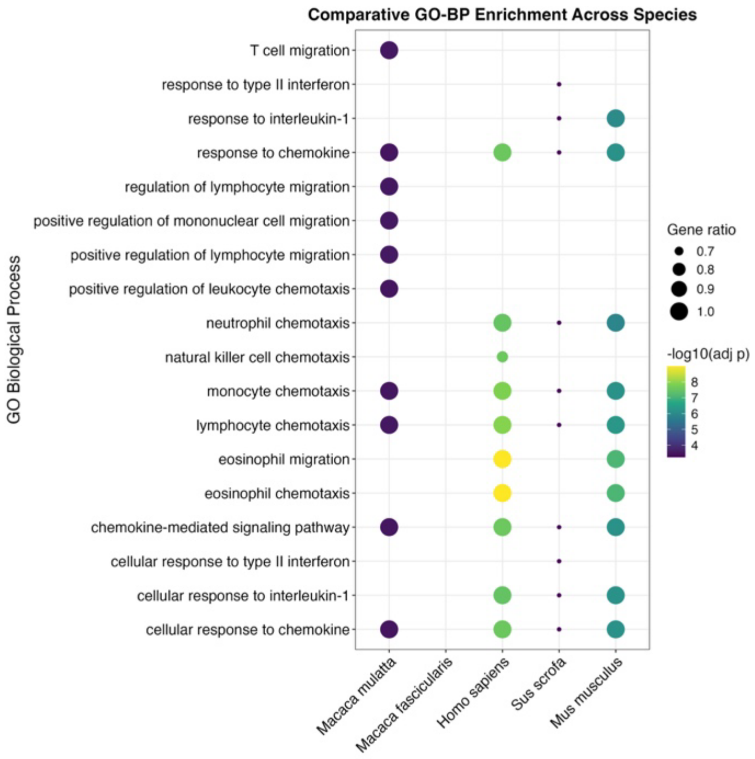


B

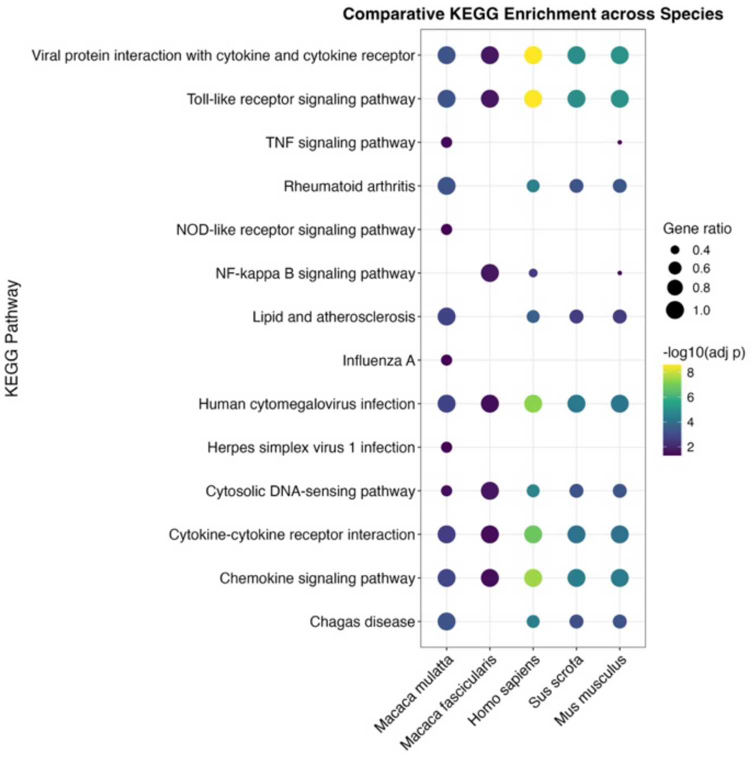


**Figure S12.** Gene Ontology term enrichment and KEGG enrichment plots for the component genes of macrogene 1490 for *Macaca mulatta, Macaca fascicularis, Homo sapiens, Sus scrofa* and *Mus musculus* in the AH atlas. (**A**) Gene ontology term enrichment plot. (**B**) KEGG enrichment plot.

**A**

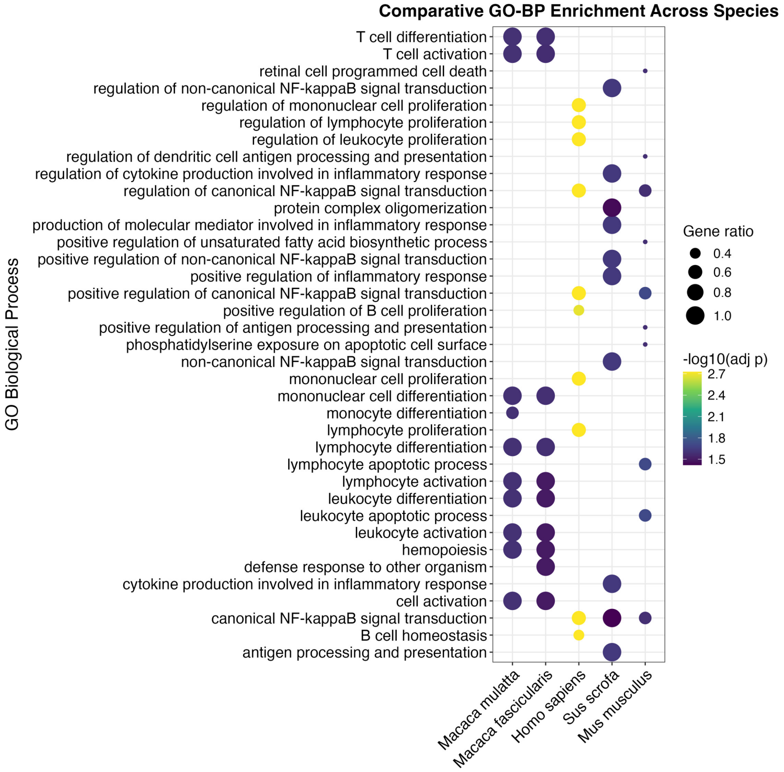


**B**

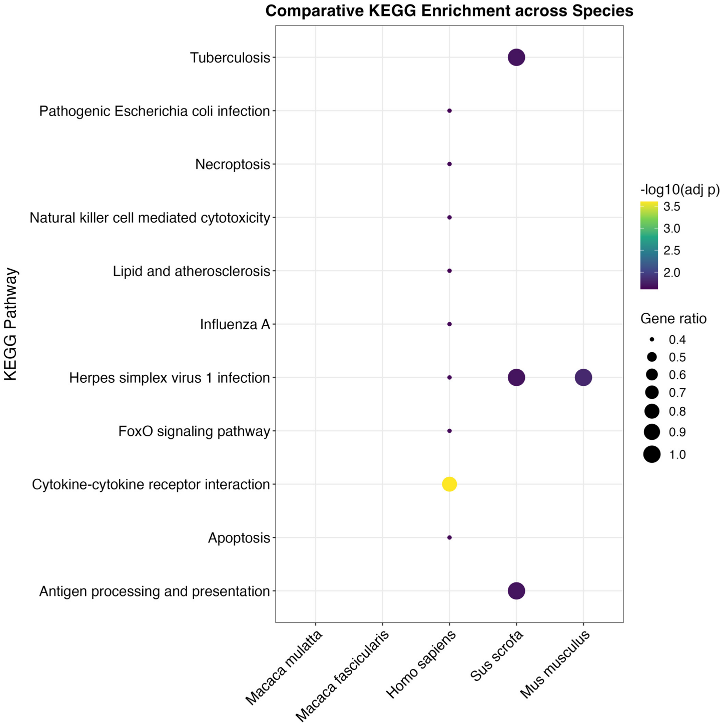


**Figure S13.** Gene Ontology term enrichment and KEGG enrichment plots for the component genes of macrogene 141 for *Macaca mulatta, Macaca fascicularis, Homo sapiens, Sus scrofa* and *Mus musculus* in the AH atlas. (**A**) Gene ontology term enrichment plot. (**B**) KEGG enrichment plot.

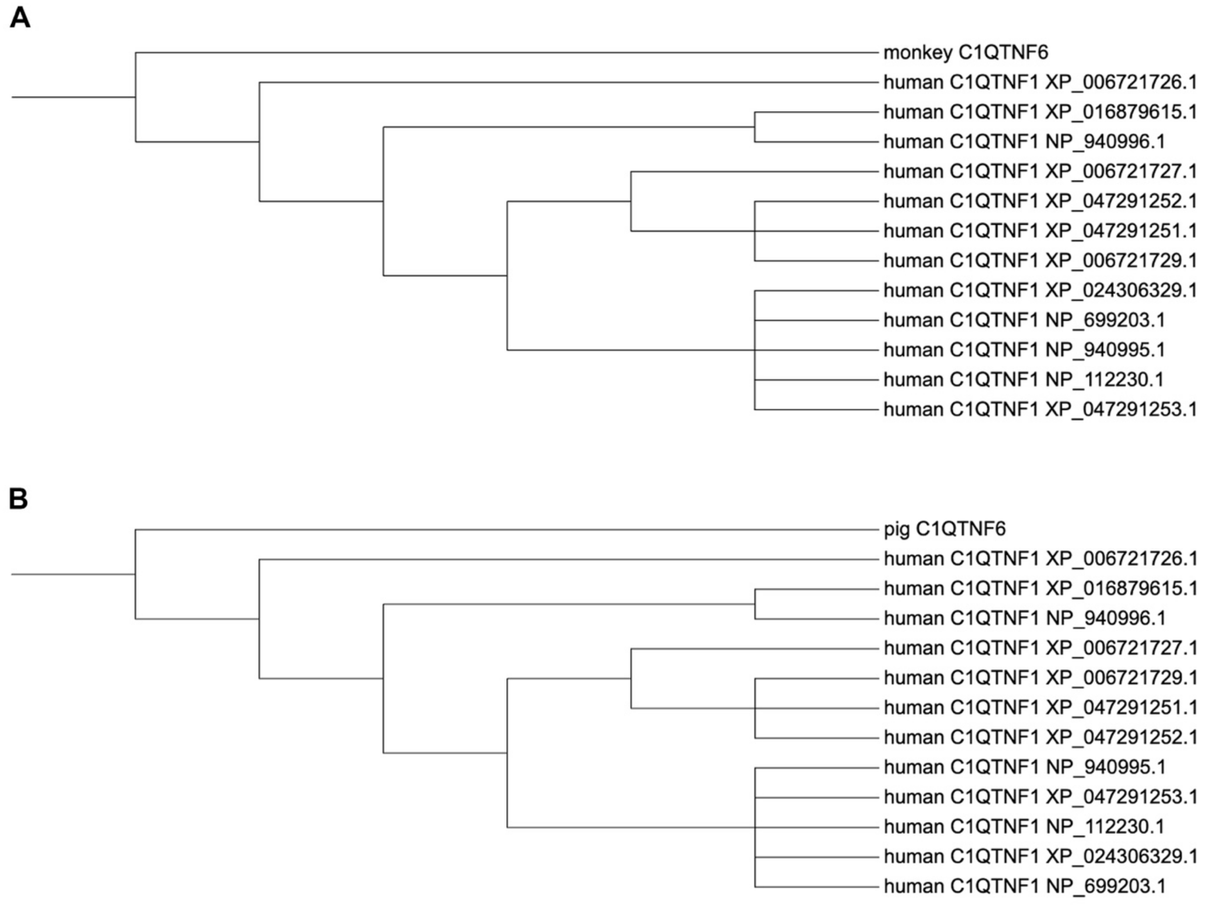


**Figure S14.** Phylogenetic tree of the C1QTNF6 proteins sourced from human (*Homo sapiens*), monkey (*Macaca mulatta*) and pig (*Sus scrofa*). (**A**) Phylogenetic relationships between C1QTNF6 protein sourced from monkey and human. (**B**) Phylogenetic relationships between C1QTNF6 protein sourced from pig and human.

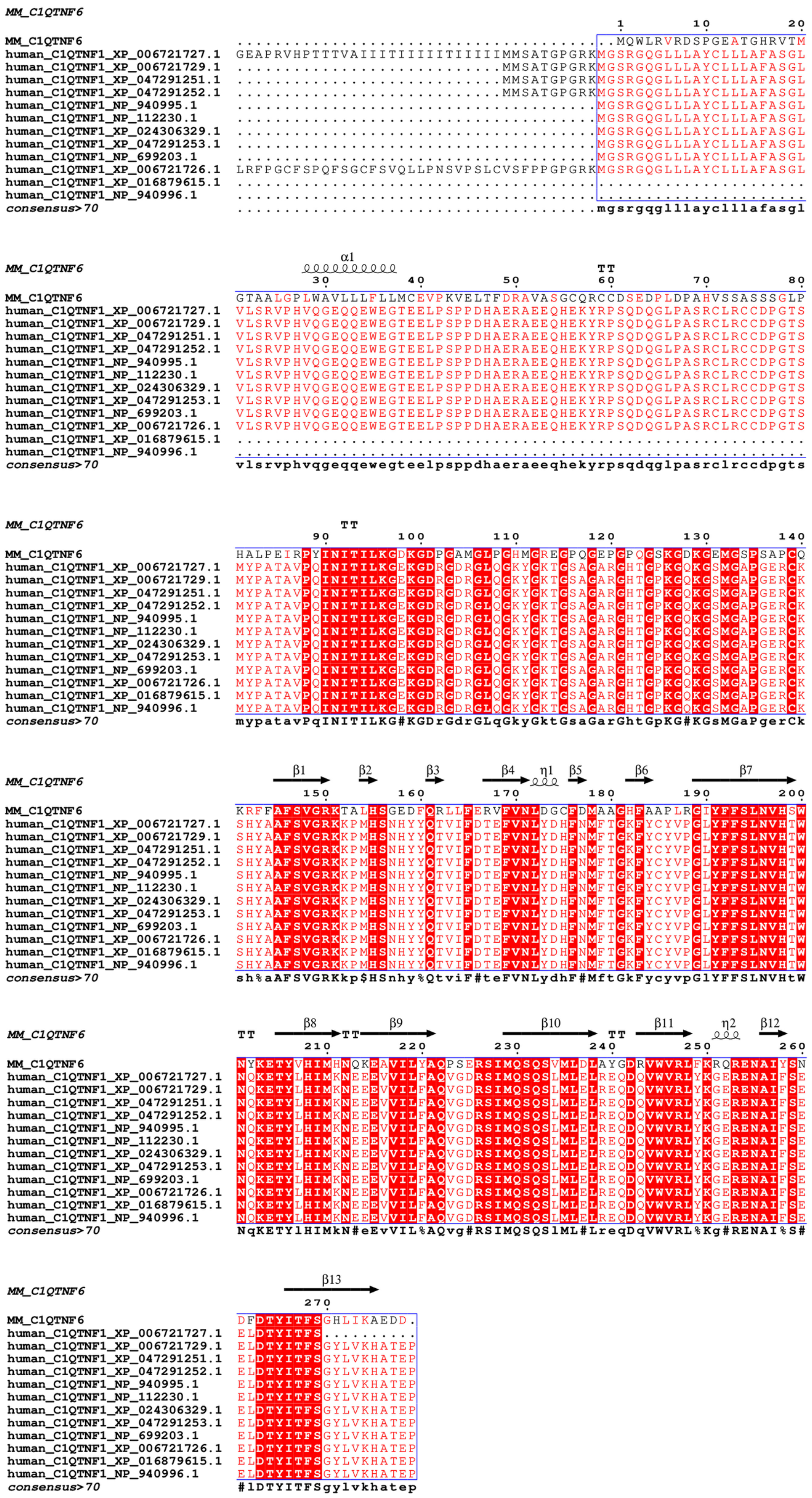


**Figure S15.** Comparison of the sequence and the structure of the C1QTNF6 proteins sourced from monkey (*Macaca mulatta*) and human (*Homo sapiens*). η: secondary structure, 310 helix. Red box with white character means strict identity. Red characters mean similarity in a group: *ISc (in-Group Score) > 0.7.* Blue frame means similarity across groups: *TSc (Total Score) > 0.7.* TT: strict β-turns.

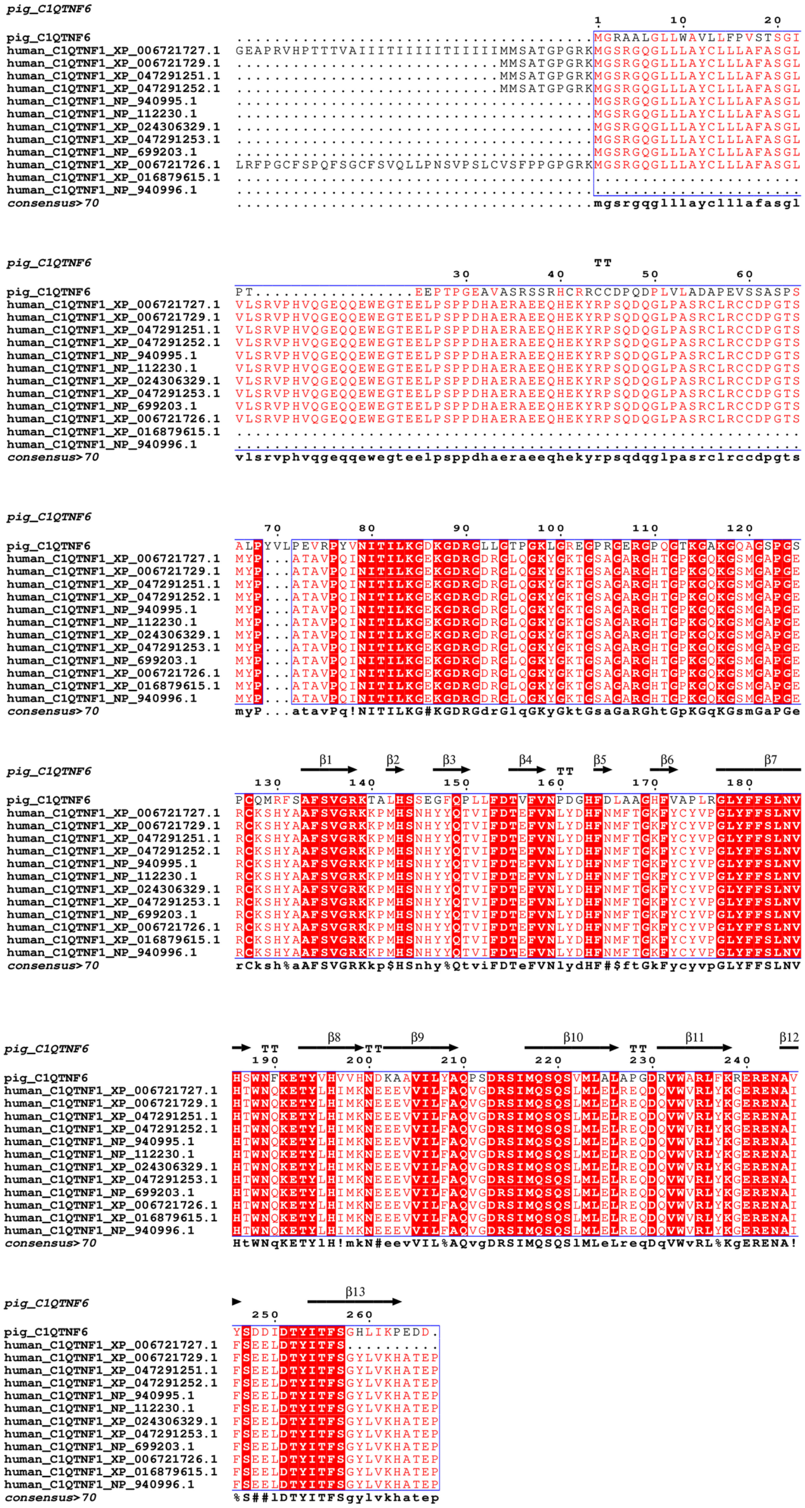


**Figure S16.** Comparison of the sequence and the structure of the C1QTNF6 proteins sourced from pig (*Sus scrofa*) and human (*Homo sapiens*). η: secondary structure, 310 helix. Red box with white character means strict identity. Red characters mean similarity in a group: *ISc (in-Group Score) > 0.7.* Blue frame means similarity across groups: *TSc (Total Score) > 0.7.* TT: strict β-turns.

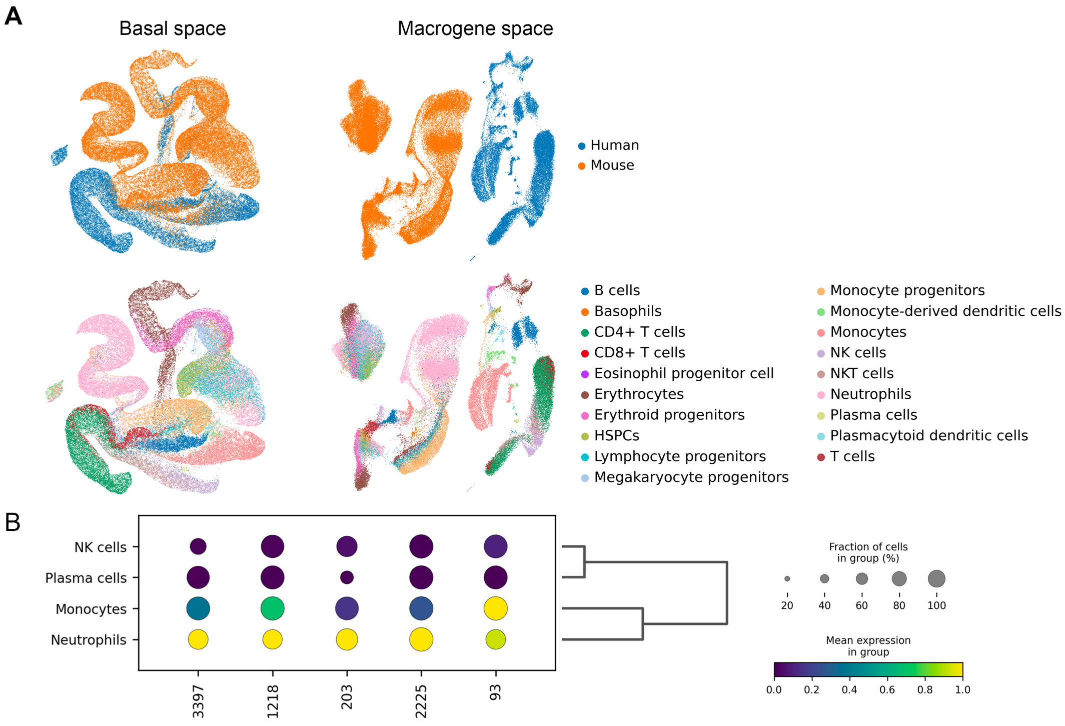


| Macrogene ID | Representative genes |
|---|---|
| 3397 | *Trp53inp1*, ***TP53INP1***, *S100a9**, *S100a8**,***S100A9*, PRDX5***, *Anxa7* |
| 1218 | *S100a9**, ***S100A9****, ***S100A12***, *S100a8**, ***S100A8*, S100A13, S100A2, RCVRN***, *Efcab9*, ***OCM*** |
| 203 | ***CAMP****, *Camp**, ***CST3***, *Cst3*, *Cst7*, ***CST7*, RARRES1***, *Ngp**, ***LXN*** |
| 2225 | ***ZP3***, *Calm3*, ***IGFBP3***, *Gna11*, ***CRB2, GAPDH, PRNP, TNFAIP6***, *Dtx3l*, ***BAG2,*** *Gnaq*, *Birc6*, *Rock1*, ***CD33****, *Ltf** |
| 93 | *S100a4**, ***S100A4****, *S100a10*, ***S100A10***, *S100a6*, ***S100A11*** |

**Purple: human genes**
Black: mouse genes
*: marker genes supported by the literature

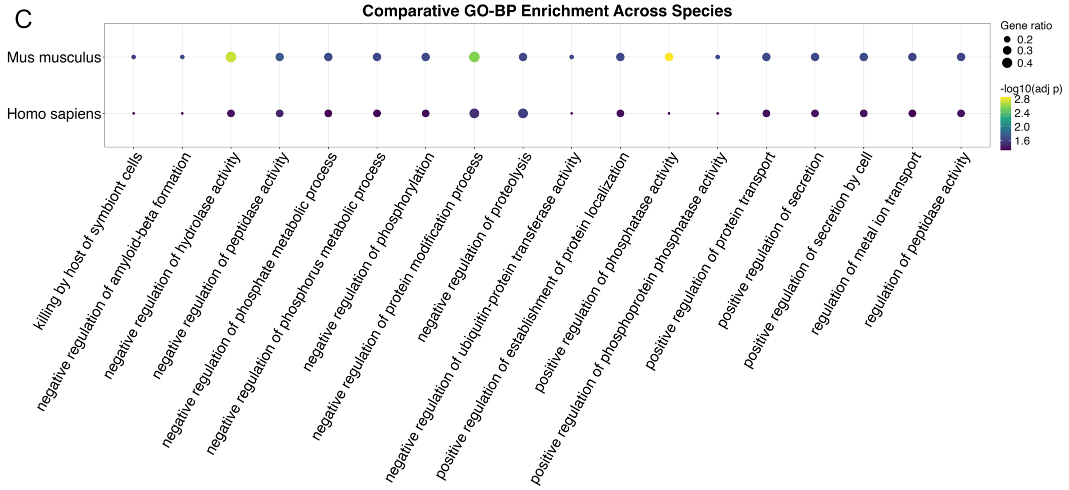


**Figure S17.** Integration results and the downstream differential macrogene analysis of immune datasets between human (*Homo sapiens*) and mouse (*Mus musculus*) using Unify. (A) UMAP plot of the macrogene space and the basal space after integration. (B) Dotplot of the expression level of the differential expressed macrogenes in macrophage (Upper) and the component genes of each differential expressed macrogenes (Lower). (C) Gene Ontology term enrichment plot of the component genes from human and mouse of macrogenes 2225.

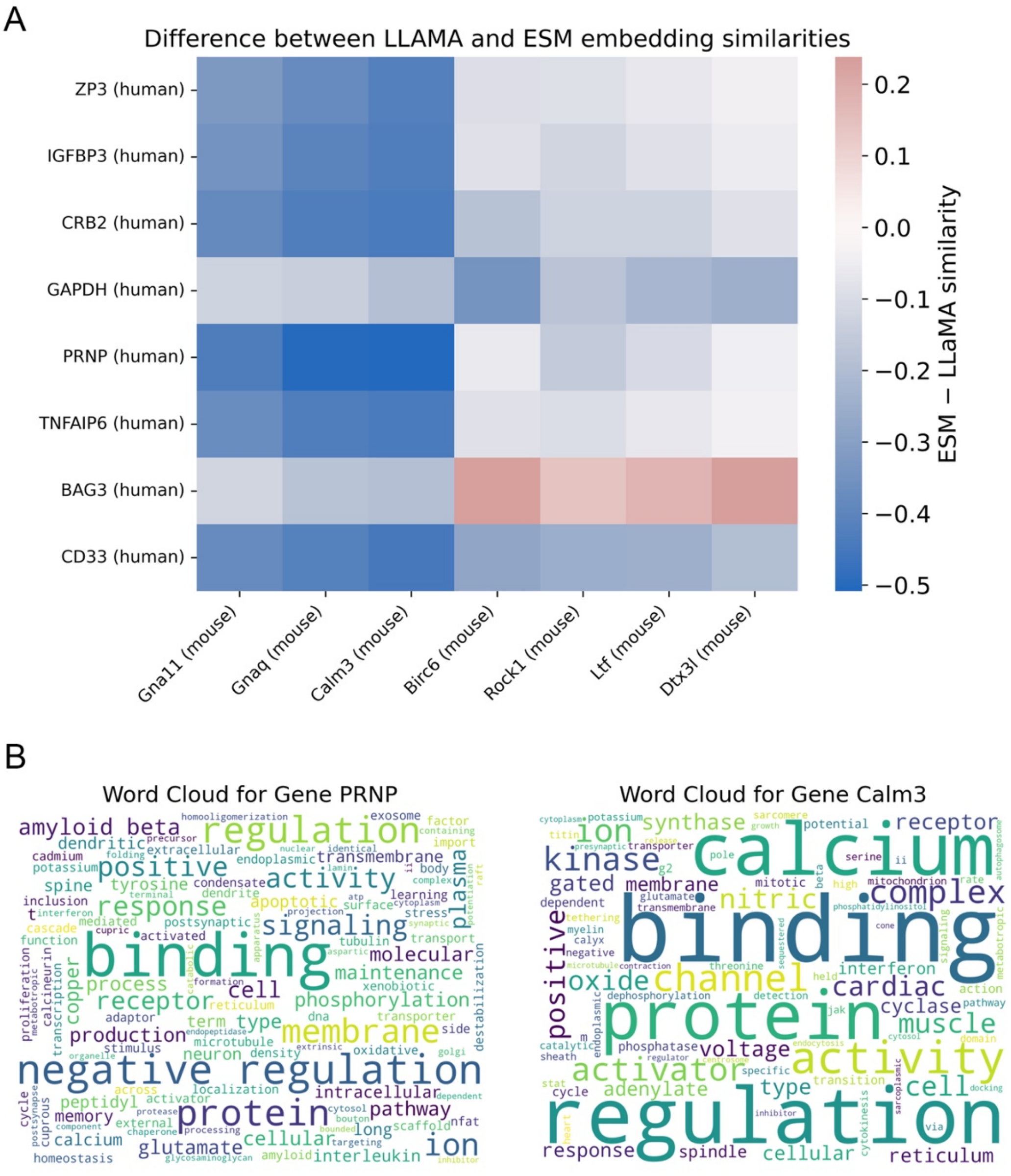


**Figure S18.** (A) Heatmap of the ESM similarity and LLAMA similarity difference of the gene pairs within the component genes of the macrogene 2225 of human and mouse. (B) Word cloud of the Gene Ontology terms description of the human gene *PRNP* and mouse gene *Calm3*.

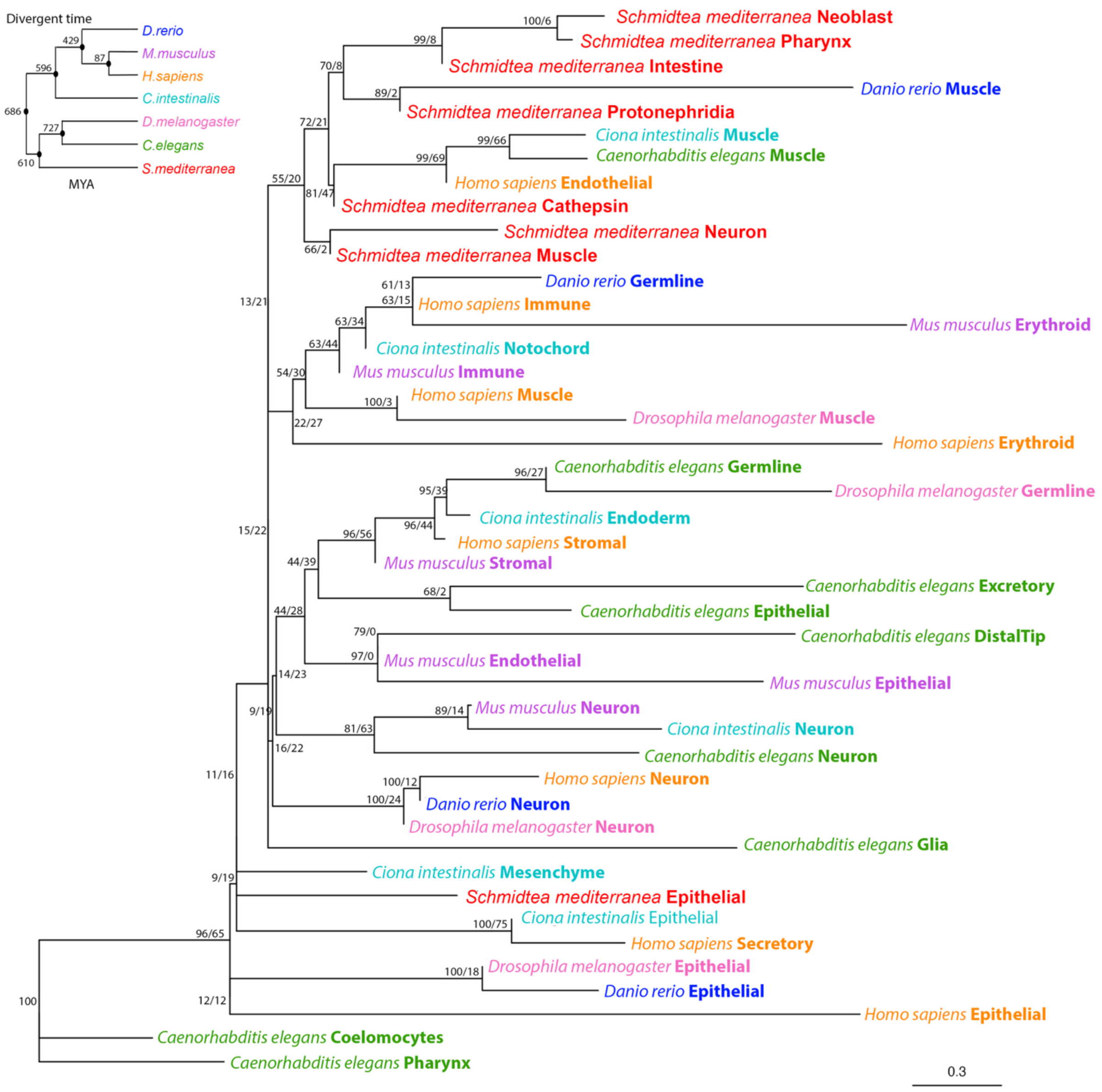


**Figure S19.** Unrooted cell type tree for seven phylogenetically distant species (*Schmidtea mediterranea, Danio rerio, Ciona intestinalis, Mus musculus, Homo sapiens, Drosophila melanogaster,* and *Caenorhabditis elegans*) constructed from the integrated basal embedding generated by SATURN, adopted from Zhong et al[1]. Species and cell types are labeled at the tips. Node support values are printed as “jumble score/scjackknife score”. MYA: million years ago.
