## Supplementary_Note for "Unify: Learning Cellular Evolution with Universal Multimodal Embeddings"

### Supplementary Note 1: Gene functional description

Functional annotations for each gene can be drawn from various sources—Gene Ontology (GO) terms, Kyoto Encyclopedia of Genes and Genomes (KEGG) pathways, Reactome pathways, or other curated descriptions. In this study, we uniformly adopted GO terms to represent gene function across all species.

For well–studied organisms (human and mouse), we retrieved the reviewed GO annotations from the UniProt database[1]. For other species, we obtained their GO annotation files directly through the Gene Ontology Consortium's download portal[2]. If GO annotations are unavailable or incomplete (in less-studied species), they can be generated de novo using orthology-based tools such as eggNOG-mapper[3, 4].

To ensure consistent and functionally meaningful gene representation across diverse species, we standardized annotations using Gene Ontology (GO) terms. Where possible, we prioritized high-confidence, manually reviewed entries (e.g., from UniProtKB/Swiss-Prot), minimizing annotation bias and maximizing cross-species comparability.

### Supplementary Note 2: Methods used in the benchmark session

#### SATURN

SATURN[5] is a weakly-supervised autoencoder that incorporates cell type information from each species individually. In combination with the protein embeddings derived from a pretrained protein language model, it is trained to reconstruct the raw count expression data using the negative log likelihood of a ZINB distribution.

#### SAMap

SAMap[6] is a method designed to integrate single-cell RNA-seq datasets across species by aligning cell types based on both gene expression and gene homology information. It extends the Self-Assembling Manifold (SAM) algorithm by incorporating cross-species gene-gene mappings to guide the integration process. SAMap constructs manifolds for each species dataset and iteratively aligns them by updating gene-gene and cell-cell correspondences, allowing it to detect conserved and species-specific cell types.

#### fastMNN

fastMNN (Fast Mutual Nearest Neighbors)[7] is a method for integrating single-cell RNA-seq datasets by identifying shared cell populations across batches. It works by detecting mutual nearest neighbors (MNNs) between batches, i.e. cells that are closest to each other in expression space despite coming from different datasets. These MNN pairs help estimate the batch effect, which is then corrected by aligning the batches in a common low-dimensional space. fastMNN improves computational efficiency by applying dimensionality reduction early and processing batches sequentially. fastMNN for cross-species data integration is restricted to orthologous genes.

#### Harmony

Harmony[8] is an algorithm designed to integrate single-cell RNA-seq datasets by iteratively adjusting the generated cell embedding to align cells from different batches. It operates in a reduced dimensional space (typically PCA), where it groups cells into clusters based on similarity and then iteratively adjusts their positions to ensure each cluster contains a balanced mix of cells from different batches. By aligning shared cell types across datasets without relying on prior annotations, Harmony effectively removes technical variation and produces corrected embeddings suitable for downstream analysis. Harmony for cross-species data integration is restricted to orthologous genes.

#### scGen

scGen[9] is a deep learning-based method that uses variational autoencoders (VAEs) to model and predict gene expression responses in single-cell RNA-seq data. It learns a latent representation of cells that captures both biological variability and batch effects. During integration, scGen disentangles these factors and uses vector arithmetic in the latent space to simulate how cells from one condition or batch would appear in another. scGen for cross-species data integration is restricted to orthologous genes.

#### scVI

scVI (single-cell Variational Inference)[10] is a deep generative model that uses variational autoencoders (VAEs) to learn a probabilistic representation of single-cell RNA-seq data. It models gene expression counts using a negative binomial distribution, accounting for both biological variation and technical noise such as batch effects. By encoding cells into a shared latent space, scVI enables effective integration across different datasets or conditions. scVI for cross-species data integration is restricted to orthologous genes.

#### BBKNN

BBKNN (Batch Balanced KNN)[11] is a method for integrating single-cell RNA-seq datasets by correcting batch effects at the level of neighborhood graph construction. BBKNN builds a graph where each cell is connected to a specified number of nearest neighbors from each batch separately. This ensures that the local neighborhood of each cell is balanced across batches, reducing technical variation while maintaining the underlying biological structure. BBKNN operates on the principal components of the datasets. BBKNN for cross-species data integration is restricted to orthologous genes.

#### Scanorama

Scanorama[12] is a tool designed to integrate single-cell RNA-seq datasets from different batches or studies. The method starts by reducing the dimensionality of each dataset, identifying mutual nearest neighbors between pairs, and then correcting differences to bring shared cell types into alignment. Subsequently, Scanorama merges the data into a unified, corrected representation. It progressively merges datasets in a way that maintains biological variation while minimizing technical noise. Scanorama for cross-species data integration is restricted to orthologous genes.

#### Seurat V4

Seurat (version 4.3.0)[13] integrates single-cell datasets from different conditions, batches, technologies, or even species by identifying shared biological signals while correcting for technical differences. After dataset normalization, highly variable gene selection and dimension reduction, Seurat uses Canonical Correlation Analysis (CCA) to project datasets into a shared space and identifies anchors, which are pairs of biologically similar cells across datasets based on the mutual nearest neighbors. Seurat further uses the anchors to align the datasets into a common coordinate system. Seurat for cross-species data integration is restricted to orthologous genes.

### Supplementary Note 3: Evaluation of Batch Correction and Biological Conservation

We treat species labels as "batches" and cell-type labels as "biological groups." After integration, we quantify batch removal and biological signal preservation for each method. For batch correction, we use the following sub-metrics: Adjusted Rand Index (ARI) batch, Adjusted Silhouette Width (ASW) batch, Normalized Mutual Information (NMI) batch, Local Inverse Simpson's Index (iLISI), kBET, graph connectivity, and Principal Component Regression. For bio-conservation, we use the following metrics: ARI cell type, ASW cell type, Cell-type LISI (cLISI), NMI cell type, highly variable gene (HVG) conservation and trajectory conservation. We run these evaluations through the scIB package[14].

#### Adjusted Rand Index (ARI)

$ARI_{batch}$ measures the overall agreement between the inferred clusters and known species labels with adjustment for chance, while the $ARI_{celltype}$ measures the overall agreement between the inferred clusters and the known cell type labels with adjustment for chance.

$$ARI = \frac{\sum_{ij}\binom{n_{ij}}{2} - [\sum_i \binom{a_i}{2} \sum_j \binom{b_j}{2}] / \binom{n}{2}}{\frac{1}{2}[\sum_i \binom{a_i}{2} + \sum_j \binom{b_j}{2}] - [\sum_i \binom{a_i}{2} \sum_j \binom{b_j}{2}] / \binom{n}{2}}$$

Where $n_{ij}$ is the number of cells in cluster i and batch/celltype j, $a_i = \sum_j n_{ij}$, $b_j = \sum_i n_{ij}$, and n is the total number of cells.

We used the Louvain clustering to compute the inferred clusters. The value ranges from 0 to 1. The higher value represents a higher match between the clusters. To make the $ARI_{batch}$ score higher, we use below equation for conversion:

$$ARI_{batch} = 1 - ARI_b$$

#### Adjusted Silhouette Width (ASW)

ASW computes the average silhouette s(i) over all cells. It measures how intermixed species/cell type labels are by comparing within-clusters and between-clusters distances, where the clusters refer to species or cell type for ASW batch and ASW cell type separately. The value ranges from -1 to 1. Higher values represent better-quality clusters.

$$s(i) = \frac{b(i) - a(i)}{max\{a(i), b(i)\}}, ASW = \frac{1}{n}\sum_{i=1}^{n} s(i)$$

Where a(i) is the average distance of cell i to cells of its own cluster, and b(i) is the minimum average distance to any other clusters.

**Graph Local Inverse Simpson's Index (LISI)**

Graph LISI quantifies the diversity in a neighborhood, ranging from 1 (minimum diversity) to B (maximum diversity), where B is the number of batches or the cell types. To standardize interpretation, LISI scores are rescaled to the interval [0,1] following Luecken et al.[14]. We first calculate the median LISI across all the cells:

$$cLISI = median\big(f(x)\big), x\ is\ each\ cell$$

$$iLISI = median\big(g(x)\big), x\ is\ each\ cell$$

We further normalize the score between 0 and 1:

For cLISI:

$$f(x) = \frac{B - x}{B - 1}$$

For iLISI:

$$g(x) = \frac{x - 1}{B - 1}$$

After normalization, higher value in cLISI and iLISI means perfect cell type separation and perfect species mixing.

**Normalized Mutual Information (NMI)**

NMI measures the similarity between two clusters. It ranges from 0 to 1. The higher value means better alignment between clusters. The NMI between cluster label C and class labels Y is defined as:

$$NMI(C, Y) = \frac{2 \times I(Y; C)}{H(C) + H(Y)}$$

where H(.) is the entropy and I(Y;C) is the mutual information between Y and C

$$I(Y; C) = \sum_{y \in Y} \sum_{c \in C} p(y, c) \log\left(\frac{p(y, c)}{p(y)p(c)}\right)$$

H(C) and H(Y) are the entropies

$$H(C) = -\sum_{c \in C} p(c) \log\ \mathrm{p(c)}, H(Y) = -\sum_{y \in Y} p(y) \log\ p(y),$$

p(c,y) is the joint probability of a cell being assigned to cluster c and label y, and p(c), p(y) are the marginal probabilities.

Similar to ARI, we use the Louvain clustering to compute the inferred clusters. For NMI batch, we reversed it by subtracting it from 1.

**kNN batch effect test (kBET)**

The kNN Batch-Effect Test (kBET) evaluates how well different batches are mixed in an integrated embedding. For each cell, kBET examines whether its local neighborhood's batch composition matches the expected global distribution. A high overall kBET value indicates successful batch mixing. We followed the code in scIB[14] to compute the kBET.

**Principal Component Regression (PCR)**

PCR quantifies how much of the variance in the top K principal components (PC) is explained by batch labels. For PC k, let $p_k = (p_{1k}, \dots, p_{nk})^T$ be the scores of all n cells along that component, and let $B_i$ be the batch label of cell i. Fit the linear model

$$p_{ik} = \beta_0 + \sum_{b=1}^{B} \beta_b 1\{B_i = b\} + \varepsilon_{ik}$$

And compute

$$R_k^2 = 1 - \frac{\sum_{i=1}^{n}(p_{ik} - \hat{p}_{ik})^2}{\sum_{i=1}^{n}(p_{ik} - \bar{p}_k)^2}$$

Where $\widehat{p_{ik}}$ are the fitted values and $\overline{p_k}$ is the mean of $p_k$.

Weight each $R_k^2$ by the variance explained $\lambda_k$ of PC k, and normalize:

$$PCR_{batch} = \frac{\sum_{k=1}^{K} \lambda_k R_k^2}{\sum_{k=1}^{K} \lambda_k}$$

Here $\lambda_k = Var(p_{1k}, \dots, p_{nk})$ is the variance explained by PC k.

**Graph Connectivity**

Graph connectivity evaluates how well cells sharing the same label (batch or cell type) remain interconnected in the integrated embedding's k-nearest-neighbor (kNN) graph. After building a kNN graph on all cells, one examines each label group by extracting the subgraph induced by that label's cells and measuring the size of its largest connected component relative to the total number of cells in the group. These proportions are then averaged over all labels to yield the overall connectivity score. High connectivity for batch labels indicates thorough mixing (cells from different batches interleave). This metric captures global structural coherence in the embedding and complements local measures (like silhouette or LISI) by assessing group-level continuity across the entire graph.

#### HVG Conservation

HVG conservation measures the extent to which integration preserves key gene-level signals that drive cellular heterogeneity. Before and after applying an integration method, we independently identify the top set of highly variable genes within each species or batch. The conservation score is then defined as the proportion of genes that remain shared between the pre-integration and post-integration HVG sets. A high HVG conservation score indicates that the integration process has maintained the most informative, biologically driven gene variation rather than smoothing it away. Conversely, a low score suggests that important gene-level differences, often linked to true cell type distinctions, have been attenuated by the integration. By focusing on gene-level variability, HVG conservation complements embedding-based metrics, ensuring that integration methods not only mix datasets effectively but also retain the critical molecular features underlying cell biology.

#### Trajectory Conservation

Trajectory Conservation (TC) quantifies how well integration preserves dynamic, pseudotemporal relationships. We infer trajectories before and after integration using Scanpy's diffusion pseudotime (sc.tl.dpt)[15] and designate the most extreme cell in each cluster as the root. The Spearman rank correlation between pre- and post-integration pseudotime values, computed via the scIB pipeline, yields the TC score, which is then normalized to [0,1]. Higher TC values indicate better preservation of biological progression through the integration process.

### Supplementary Note 4: Cell type tree construction

We built cross-species cell-type phylogenies following Jasmine *et al*[16]. First, for both SATURN and Unify embeddings, we performed PCA and retained the top 20 components to reduce noise and computational load. To represent each of the 45 annotated cell types, we randomly sample one cell per type, yielding a 45 × 20 data matrix.

Phylogenetic inference was carried out using PHYLIP's contml (v3.698) under a Brownian motion model of continuous character evolution. We enabled the continuous-data option (–C) and global rearrangements (–G), and randomized taxon order 100 times per run (–J 100), selecting the highest-likelihood tree each time. We repeated this process for 250 independent runs with 100 jitters per run, then midpoint-rooted the best-scoring tree for presentation.

To evaluate tree robustness, we calculate two repeatability measures:

**(1)** Technical repeatability (jumble score): the consistency of topology across random taxon-order perturbations.
**(2)** Biological repeatability (sc-jackknife score): the recovery rate of the original tree when resampling one cell per type (500 replicates).

Together, these analyses quantify both the stability and the biological fidelity of our integrated cell-type trees.
